# The interferon-stimulated gene product HERC5 inhibits human LINE-1 retrotransposition with an ISGylation-independent mechanism

**DOI:** 10.1101/2025.09.09.675047

**Authors:** Kei Nishimori, Ahmad Luqman-Fatah, Yuzo Watanabe, Mari Takahashi, Takuhiro Ito, Fuyuki Ishikawa, Tomoichiro Miyoshi

**Affiliations:** Laboratory for Retrotransposon Dynamics, RIKEN Center for Integrative Medical Sciences, Yokohama 230-0045, Japan; Department of Gene Mechanisms, Graduate School of Biostudies, Kyoto University, Kyoto 606-8501, Japan; Proteomics Facility, Graduate School of Biostudies, Kyoto University, Kyoto 606-8502, Japan; Laboratory for Translation Structural Biology, RIKEN Center for Integrative Medical Sciences, Yokohama 230-0045, Japan; Graduate School of Medical and Dental Sciences, Institute of Science Tokyo, Tokyo 113-8510, Japan

**Author notes:** Corresponding author: Please address correspondence to Tomoichiro Miyoshi.

## Abstract

Mobilization of long interspersed element-1 (LINE-1 or L1) compromises genome stability and can cause sporadic genetic disease. Accordingly, cells have evolved multiple mechanisms to restrict L1 retrotransposition. Several interferon-stimulated genes (ISGs) that interact with cytoplasmic L1 ribonucleoproteins (RNPs), which contain the L1-encoded proteins ORF1p and ORF2p, have been identified as suppressors of L1 retrotransposition. We previously reported that the ISG protein HECT and RLD domain containing E3 ubiquitin-protein ligase 5 (HERC5) efficiently inhibits L1 retrotransposition. While HERC5 is known to restrict numerous viruses through ISGylation, how HERC5 inhibits L1 remains to be elucidated. Here, we show that HERC5 inhibits L1 retrotransposition through an ISGylation-independent mechanism. HERC5 interacts with L1 RNA and reduces ORF1p levels, a function that requires the *ORF1* coding sequence. We further demonstrate that HERC5 decreases L1 translation efficiency and prevents the assembly of L1 RNPs. Our comparative analysis suggests that HERC5 may have acquired its L1-inhibitory function during the evolution of the small HERC family. These findings uncover a previously unidentified mechanism by which an ISG protein recognizes and inhibits L1 and suggest a role for HERC5 as an evolutionarily adapted restriction factor that expands the repertoire of cellular defenses against retrotransposons.

## Introduction

Human genome sequencing revealed that the protein-coding regions cover only a small portion of the genome at less than 2% (1, 2). In comparison, nearly half of the genome is made up of transposable elements (1–3). Among them, Long INterspersed Element-1 (LINE-1 or L1) comprises ∼17% of the human genome (1, 2). There are more than 500,000 L1 copies in the human genome; however, only ∼80–100 full-length L1 copies are estimated to be retrotransposition-competent L1s (RC-L1s) (4–6).

RC-L1s are ∼6 kb in length and contain a 5′ untranslated region (UTR), two open reading frames (*ORF1* and *ORF2*), and a 3′ UTR (3, 7). L1s are transcribed from an internal RNA polymerase II promoter within the 5′ UTR (8–11). *ORF1* encodes a ∼40 kDa protein (ORF1p) with RNA-binding and nucleic acid chaperone activities (12, 13). ORF1p forms a homotrimeric structure and is essential for L1 retrotransposition (14, 15). *ORF2* encodes a ∼150 kDa protein (ORF2p) possessing endonuclease (EN) and reverse transcriptase (RT) activities, both of which are required for L1 retrotransposition (16–19). In the cytoplasm, ORF1p and ORF2p preferentially bind to L1 RNA in cis and form L1 RiboNucleoProtein (RNP) complexes (20–23). L1 RNPs then gain nuclear access; however, the mechanism remains unclear (24). In the nucleus, ORF2p EN activity introduces a single-strand endonucleolytic cleavage at 5′-TTTTT/AA-3′ (the slash indicates the ORF2p cleavage site) and related variants of that consensus sequence in genomic DNA, exposing a 3′-OH group (16, 25, 26). The T-rich sequence, with the liberated 3′-OH group, base-pairs with the L1 RNA poly(A) tail and provides a primer for reverse transcription of L1 cDNA from L1 RNA (22, 27). Subsequently, L1 cDNA is integrated into the genome through a process called target-site primed reverse transcription (TPRT) (28–30).

*De novo* retrotransposon insertions can disrupt genes or alter gene expression, potentially leading to sporadic human diseases (31, 32). To date, more than 100 L1-mediated retrotransposition events have been implicated in disease-causing mutations (33, 34), including L1 insertion into *F8* (haemophilia A), *APC* (colon cancer), and SVA insertion into *FKTN* (Fukuyama congenital muscular dystrophy) (35–37). In addition to insertional mutagenesis, recent studies have suggested that retrotransposon intermediates, including L1 cDNA, L1 RNA, and EN-induced DNA breaks, can contribute to the development of autoimmune diseases, cancer progression, and cellular senescence (38–44). To maintain genomic stability, host cells have evolved multiple defense mechanisms to suppress L1 retrotransposition and its expression (e.g., Apolipoprotein B mRNA-editing enzyme catalytic polypeptide 3 [APOBEC3], zinc-finger antiviral protein [ZAP or ZC3HAV1], Moloney leukemia virus type 10 protein [MOV10]) (45–51). Elucidating the molecular mechanisms by which host factors regulate potentially pathogenic L1 activity is expected to provide insights into future preventive medicine and novel therapeutic strategies for multiple diseases.

To identify the host proteins involved in the regulation of L1 retrotransposition, we previously performed immunoprecipitation of wild-type ORF1p and an RNA-binding mutant (RBM) of ORF1p, followed by liquid chromatography-tandem mass spectrometry (LC–MS/MS) and label-free quantification (LFQ) analysis (52). We identified helicase with zinc finger 2 (HELZ2), HECT and RLD domain containing E3 ubiquitin-protein ligase 5 (HERC5), and 2′-5′-oligoadenylate synthetase-like (OASL) as ORF1p interactors, and overexpression of these factors significantly inhibited L1 retrotransposition (52). HELZ2 suppresses L1 retrotransposition by recognizing the L1 5′ UTR and reducing L1 RNA levels (52). However, the molecular mechanism by which HERC5 inhibits L1 remained unclear.

HERC5, originally identified as cyclin E-binding protein 1 (Ceb1), is a member of the HERC family, which is characterized by the presence of a homologous to the E6-AP carboxyl terminus (HECT) domain and one or more RCC1-like domains (RLDs) (53–55). HERC5 is an interferon-stimulated gene (ISG) product and a major E3 ligase for ISGylation (56). ISGylation is a ubiquitin-like post-translational modification in which ISG15, a ubiquitin-like 15 kDa protein containing two ubiquitin-like domains, is covalently conjugated to target substrates through an E1 activating enzyme (UBA7/UBE1L), an E2 conjugating enzyme (UBE2L6/UbcH8), and E3 ligases (HERC5, TRIM25, and ARIH1), similar to ubiquitin modification (56–62). HERC5 is a ∼116 kDa protein and the HECT domain, especially the catalytic cysteine at position 994, is essential for ISGylation (56, 63). Additionally, the RLD domain interacts with polysomes, facilitating co-translational ISGylation of newly synthesized viral and host proteins, thereby playing an important role in innate immunity (64, 65). HERC5 inhibits various viruses, including influenza A virus (IAV), human immunodeficiency virus type 1 (HIV-1), and Ebola virus through an ISGylation-dependent mechanism (66–68). Although an ISGylation-independent mechanism has also been reported for the inhibition of HIV-1 and Ebola virus (69, 70), the detailed mechanism remains largely unknown.

In this study, we elucidated the molecular mechanism by which HERC5 inhibits L1 retrotransposition. Our results revealed that HERC5 suppresses L1 retrotransposition independently of ISGylation. We also found that the HERC5 RLD domain binds to L1 RNPs via RNA, leading to a decrease in L1 translation efficiency and inhibition of L1 RNP formation. Finally, comparative analysis across the small HERC paralogs suggests that HERC5 acquired the ability to inhibit L1 during the evolution of the small HERC family.

## Materials and Methods

### Cell lines and cell culture conditions

HEK293T and HeLa-JVM (19, 71, 72) cell lines were grown in Dulbecco’s Modified Eagle Medium (DMEM) (Nissui, Tokyo, Japan or Shimadzu Diagnostics, Tokyo, Japan) supplemented with 10% (volume/volume [v/v]) fetal bovine serum (FBS) (Gibco, Amarillo, Texas, United States; Capricorn Scientific, Ebsdorfergrund, Germany; or MP Biomedicals, California, United States), 0.165% (weight/volume [w/v]) NaHCO_3_ (Nacalai Tesque, Kyoto, Japan), 100 U/mL penicillin G (Sigma-Aldrich, St. Louis, MO, United States), 100 µg/mL streptomycin (Sigma-Aldrich), and 2 mM L-glutamine (Sigma-Aldrich). HeLa-HA (71, 72) cells were grown in Minimum Essential Medium (MEM) (Gibco) supplemented with 10% (v/v) fetal bovine serum (FBS), 0.165% (w/v) NaHCO_3_, 100 U/mL penicillin G, 100 µg/mL streptomycin, 2 mM L-glutamine, and 1×MEM Non-Essential Amino Acids Solution (Nacalai Tesque). In the retrotransposition assay for mouse TG_F_21, ORFeus-Mm, and zebrafish ZfL2-2, HEK293T cells were grown in Dulbecco’s Modified Eagle Medium (DMEM) high glucose medium (Nacalai Tesque) supplemented with 10% (v/v) fetal bovine serum (FBS) (Gibco), 0.165% (w/v) NaHCO_3_, 100 U/mL penicillin G, 100 µg/mL streptomycin, and 2 mM L-glutamine. The cell lines were grown at 37°C in 100% humidified incubators supplied with 5% CO_2_. The absence of Mycoplasma spp. was confirmed with VenorGeM Classic Mycoplasma Detection Kit (Sigma-Aldrich). STR genotyping was performed to confirm the identity of HeLa-JVM, HeLa-HA, and HEK293T cells by BEX (Tokyo, Japan).

### Plasmids used in this study

Plasmids for mammalian transfection experiments were purified using the Midiprep Plasmid DNA Kit (QIAGEN, Hilden, Germany) or the GenElute HP Plasmid Miniprep Kit (Sigma-Aldrich). All L1-expressing plasmids contain a retrotransposition-competent L1 (L1.3, GenBank: L19088 (4)) and have been subcloned into the pCEP4 vector backbone (Invitrogen, Massachusetts, United States) unless otherwise noted. The amino acid residues of ORF1p or ORF2p were counted from the first methionine of L1.3 ORF1p and ORF2p, respectively. The plasmids used in this study are listed below:

<u>pCEP4 (Invitrogen)</u>: is a vector backbone for mammalian expression. This plasmid was used for cloning pJM101/L1.3 and pJJ101/L1.3 variants.

<u>phrGFP-C (Agilent Technologies, California, United States)</u>: was previously described in (27). This plasmid contains a humanized Renilla green fluorescent protein gene, which is driven by the CMV promoter.

<u>pMD2.G</u>: is a lentivirus envelope expression vector that was generously provided by Dr. Didier Trono (Addgene plasmid #12259).

<u>psPAX2</u>: is a lentivirus packaging vector that was generously provided by Dr. Didier Trono (Addgene plasmid #12260).

<u>pCMV-3Tag-9 (Agilent Technologies)</u>: is a vector backbone for mammalian expression. This plasmid fuses three copies of *MYC* epitope tags to the C-terminus of proteins of interest. The protein expression is driven by a cytomegalovirus immediate early (CMV) promoter and has a hygromycin resistance gene as a selectable marker.

<u>pEBNA</u>: is a vector backbone for mammalian expression and was previously described in (73). The protein expression was driven by the EF1α promoter and has a puromycin resistance gene as a selectable marker.

<u>pcDNA6</u>: was previously described in (74). It is a derivative of pcDNA6/TR (Invitrogen) and expresses the blasticidin-resistant gene but lacks the *TetR* gene, generated by Dr. John B. Moldovan (University of Michigan).

<u>pJM101/L1.3</u>: was previously described in (4, 19). This plasmid contains the full-length retrotransposition-competent L1.3 (4), cloned into the pCEP4 vector. L1 expression is driven by both the CMV and L1.3 5′ UTR promoters. The *mneoI* retrotransposition indicator cassette was inserted into the L1.3 3′ UTR as described previously (19).

<u>pJM101/L1.3FLAG</u>: was previously described in (47). This plasmid is derived from pJM101/L1.3 and contains a single *FLAG* epitope tag sequence fused in frame to the 3′ end of the L1.3 *ORF1*-coding sequence.

<u>pJM105/L1.3</u>: was previously described in (20). This plasmid is derived from pJM101/L1.3, but contains the D702A mutation in *ORF2*, therefore expressing reverse transcriptase-deficient ORF2p, which cannot support retrotransposition.

<u>pJJ101/L1.3</u>: was previously described in (75). This plasmid is similar to pJM101/L1.3 except that it contains an *mblastI* retrotransposition indicator cassette within the L1 3′ UTR.

<u>pJJ105/L1.3</u>: was previously described in (75). This plasmid is derived from pJJ101/L1.3, but contains the D702A mutation in *ORF2*.

<u>pTMF3</u>: was previously described in (74). This plasmid is derived from pJM101/L1.3. A single *T7 gene 10* epitope tag was fused in frame to the 3′ end of the *ORF1* sequence. Three *FLAG* epitope tags were fused in frame to the 3′ end of the *ORF2* sequence.

<u>pTMH3</u>: is derived from pTMF3, but a single *hemagglutinin* epitope tag sequence is fused in frame to the 3′ end of the *ORF2* sequence, instead of the *FLAG* epitope tag.

<u>pTMF3_Δ5UTR</u>: was previously described in (52). This plasmid is derived from pTMF3, but the L1.3 5′ UTR is deleted.

<u>pL1(5&3UTRs)_Fluc:</u> was previously described in (52). This plasmid is derived from pTMF3, but the L1.3-coding region was replaced with a firefly luciferase gene.

<u>cep99-gfp-L1.3</u>: was previously described in (52, 74). This plasmid contains the full-length L1.3 with an enhanced green fluorescent protein retrotransposition indicator cassette (*mEGFPI*) (76) in the L1.3 3′ UTR. The L1.3 was cloned into a modified-pCEP4 vector that contains a puromycin-resistant gene as a selectable marker. L1.3 expression is driven by the L1 5′ UTR promoter.

<u>cep99-gfp-L1.3RT</u>(-): was previously described in (74). This plasmid is similar to cep99-gfp-L1.3, but contains the D702A mutation in *ORF2*.

<u>cep99-gfp-L1.3RT</u>(-)<u>intronless</u>: was previously described in (52, 74). This plasmid is similar to cep99-gfp-L1.3RT(-), but lacks the intron sequence in the *mEGFPI* cassette, resulting in EGFP expression independent of retrotransposition.

<u>cepB-gfp-L1.3</u>: was previously described in (52, 74). This plasmid is similar to cep99-gfp-L1.3, but contains the blasticidin S-deaminase gene instead of the puromycin-resistant gene as a selectable marker.

<u>cepB-gfp-L1.3RT</u>(-): was previously described in (74). This plasmid is similar to cepB-gfp-L1.3, but contains the D702A mutation in *ORF2*.

<u>cepB-gfp-L1.3RT</u>(-)<u>intronless</u>: was previously described in (52, 74). This plasmid is similar to cepB-gfp-L1.3RT(-), but lacks the intron sequence in the *mEGFPI* cassette, resulting in EGFP expression independent of retrotransposition.

<u>cep99-gfp-Z2</u>: was previously described in (77). This plasmid contains the full-length active zebrafish L2-2 (ZfL2-2) (78) with the *mEGFPI* cassette in the 3′ UTR of ZfL2-2. The ZfL2-2 has been cloned into the modified-pCEP4 vector that contains a puromycin-resistant gene as a selectable marker.

<u>cep99-gfp-Z2-RTm</u>: was previously described in (77). This plasmid is similar to cep99-gfp-Z2, but contains the D689Y mutation in the RT domain, which abolishes ZfL2-2 retrotransposition.

<u>cep99-gfp-ORFeus-Mm</u>: was previously described in (77) as cep99-gfp-L1SM. This plasmid contains a full-length synthetic mouse L1 element and the *mEGFPI* cassette in the 3′ UTR of the L1 element (79). The synthetic mouse L1 element named ORFeus-Mm has been cloned into the modified-pCEP4 vector that contains a puromycin-resistant gene as a selectable marker.

<u>cep99-gfp-ORFeus-Mm mut2</u>: was previously described in (77) as cep99-gfp-L1SMmut2. This plasmid is similar to cep99-gfp-ORFeus-Mm, but contains D212G and D709Y mutations in *ORF2*, which inactivate the endonuclease and reverse transcriptase activities, respectively.

<u>cep99-gfp-TG_F_21</u>: was previously described in (77). This plasmid contains a full-length mouse TG_F_21 L1 element (80) with the *mEGFPI* cassette in the 3′ UTR of the TG_F_21 L1 element (79). TG_F_21 was cloned into the modified-pCEP4 vector that contains a puromycin-resistant gene as a selectable marker.

<u>pALAF008_L1.3FLAG_M8 (RBM)</u>: was previously described in (52). This plasmid is derived from pJM101/L1.3FLAG and contains the R206A, R210A, and R211A mutations in *ORF1*, resulting in severely impaired RNA-binding activity of ORF1p (15).

<u>pALAF015_hHELZ2L-3×MYC</u>: was previously described in (52). Human *HELZ2* cDNA was cloned into pCMV-3Tag-9, enabling its expression with three copies of a MYC tag at the C-terminus driven by the CMV promoter.

<u>pALAF023_hHERC5-3×MYC</u>: was previously described in (52). Human *HERC5* cDNA was cloned into pCMV-3Tag-9, enabling its expression with three copies of a MYC tag at the C-terminus driven by the CMV promoter.

<u>pALAF024_hMOV10-3×MYC</u>: was previously described in (52). Human *MOV10* cDNA was cloned into pCMV-3Tag-9, enabling its expression with three copies of a MYC tag at the C-terminus driven by the CMV promoter.

<u>pLKO.1-TurboGFP</u>: is a lentivirus vector (Sigma Aldrich: SHC002) for expression of a *TurboGFP-*targeting shRNA sequence, containing a puromycin resistance gene as a selectable marker. This shRNA construct does not target the *EGFP* gene due to sequence differences. Cells expressing this shRNA construct were used as a shControl for the shRNA-based experiments described in this study.

<u>pALAF034_shHERC5_3_pLKO.1</u>: is a lentivirus vector that expresses shRNA targeting the coding sequence (5′-GAAGGACTAGACAATCAGAAA-3′) of human *HERC5*.

<u>pALAF060_hHERC5-3×MYC_C994A</u>: This plasmid is similar to pALAF023_hHERC5-3×MYC, but contains the C994A mutation, therefore expressing ISGylation activity-deficient HERC5.

<u>pALAF061_hHERC5-3×MYC_ΔHECT</u>: This plasmid is similar to pALAF023_hHERC5-3×MYC, but lacks the HECT domain.

<u>pALAF062_hHERC5-3×MYC_ΔRLD</u>: This plasmid is similar to pALAF023_hHERC5-3×MYC, but lacks the RLD domain.

<u>pTM489_pTMO2F3_*Alu*</u>: was previously described in (52) as pTMO2F3_*Alu*. This plasmid co-expresses *Alu* RNA and monocistronic L1.3 ORF2p with three FLAG epitope tags at the C-terminus of ORF2p. This plasmid contains the *AluY* with a neomycin resistance gene as the retrotransposition indicator cassette (81). L1.3 *ORF2* and *AluY* expressions were augmented by the CMV promoter and the 7SL enhancer, respectively. This arrangement enables the quantification of the *Alu* retrotransposition efficiency by counting the number of G418-resistant foci.

<u>pKO001_pTMO2F3D145AD702A_*Alu*</u>: was previously described in (52). This plasmid is similar to pTM489_pTMO2F3_*Alu*, but contains D145A and D702A mutations in *ORF2*, which inactivate the endonuclease and reverse transcriptase activities, respectively. These mutations render the *Alu* retrotransposition-incompetent.

<u>pKN040_pTMF3_*Alu*</u>: This plasmid is similar to pTM489 but co-expresses *Alu* RNA and the full-length bicistronic L1.3. The *T7 gene 10* and three *FLAG* epitope tags were fused in frame to the 3′ end of the *ORF1* and *ORF2* sequences, respectively. This plasmid contains the *AluY* with a neomycin resistance gene as a retrotransposition indicator cassette (81). L1.3 and *AluY* expressions were augmented by the CMV promoter and the 7SL enhancer, respectively.

<u>pKN041_pTMF3D702A_*Alu*</u>: This plasmid is similar to pKN040, but contains the D702A mutation in *ORF2*, which inactivates the reverse transcriptase activity. This mutation renders the *Alu* element retrotransposition-incompetent.

<u>pTMO2F3</u>: was previously described in (27, 74). This plasmid expresses monocistronic L1.3 ORF2p with three FLAG epitope tags at the C-terminus of ORF2p. A CMV promoter and the L1 5′ UTR augment the L1.3 ORF2p expression.

<u>pKN035_pTMO1T7</u>: expresses monocistronic L1.3 ORF1p with the T7 gene 10 epitope tag at the C-terminus of ORF1p. A CMV promoter and the L1 5′ UTR augment the L1.3 ORF1p expression.

<u>pCMV-3Tag-8-Barr</u>: was previously described in (52). Human *ARRB2* cDNA was cloned into pCMV-3Tag-8 (Agilent Technologies), enabling its expression with three copies of a FLAG tag at the C-terminus driven by the CMV promoter.

<u>pKN002_hHERC5-3×FLAG</u>: Human *HERC5* cDNA was cloned into pCMV-3Tag-8, enabling its expression with three copies of a FLAG tag at the C-terminus driven by the CMV promoter.

<u>pKN003_hHERC5-3×FLAG_ΔRLD</u>: This plasmid is similar to pKN002_hHERC5-3×FLAG, but lacks the RLD domain.

<u>pKN004_hHERC5-3×FLAG_ΔHECT</u>: This plasmid is similar to pKN002_hHERC5-3×FLAG, but lacks the HECT domain.

<u>pKN005_hHERC5-3×FLAG_C994A</u>: This plasmid is similar to pKN002_hHERC5-3×FLAG, but contains the C994A mutation, therefore expressing ISGylation-deficient HERC5.

<u>pKN024_pJM101ORFeus</u>: This plasmid is derived from pJM101/L1.3, but the *ORF1-* and *ORF2*-coding sequences were replaced with the codon-optimized human L1 *ORFeus* sequence from pDA093 (43), which was generously provided by Dr. Kathleen Burns (Addgene plasmid #131390).

<u>pKN025_hHERC5-3×MYC_pEBNA</u>: Human *HERC5* cDNA fused with three copies of a MYC tag sequence in frame to the 3′ end was cloned into pEBNA, enabling its expression driven by the EF1α promoter.

<u>pKN026_hHERC5-3×MYC_ΔRLD_pEBNA</u>: This plasmid is similar to pKN025_hHERC5-3×MYC_pEBNA, but lacks the RLD domain.

<u>pKN027_hHERC5-3×MYC_ΔHECT_pEBNA</u>: This plasmid is similar to pKN025_hHERC5-3×MYC_pEBNA, but lacks the HECT domain.

<u>pKN028_hHERC5-3×MYC_C994A_pEBNA</u>: This plasmid is similar to pKN025_hHERC5-3×MYC_pEBNA, but contains the C994A mutation, therefore expressing ISGylation activity-deficient HERC5.

<u>pKN031_hHERC4-3×MYC</u>: Human *HERC4* cDNA was cloned into pCMV-3Tag-9, enabling its expression with three copies of a MYC tag at the C-terminus driven by the CMV promoter.

<u>pKN032_hHERC6-3×MYC</u>: Human *HERC6* cDNA was cloned into pCMV-3Tag-9, enabling its expression with three copies of a MYC tag at the C-terminus driven by the CMV promoter.

<u>pVan583</u>: was previously described in (82, 83). This plasmid is derived from pJM101/L1.3, where *EGFP* and *mCherry* were conjugated to *ORF1* and *ORF2* to the 3′ termini, respectively. This plasmid was generously provided by Dr. Zbigniew Warkocki.

<u>pKN033_pVan583_EGFP</u>: is derived from pVan583. The L1 5′ UTR, *ORF1*, and *mCherry*-fused *ORF2* sequences in pVan583 were replaced with *EGFP*, allowing the *EGFP* expression alone by the CMV promoter.

<u>pKN036_mHERC6_3×MYC</u>: Mouse *HERC6* cDNA was cloned into pCMV-3Tag-9, enabling its expression with three copies of a MYC tag at the C-terminus driven by the CMV promoter.

<u>pKN039_cepB-gfp-L1 ORFeus</u>: This plasmid is similar to cepB-gfp-L1.3, but the *ORF1-* and *ORF2-*coding sequences were replaced with the codon-optimized human L1 *ORFeus* sequence from pDA093 (43).

### Virus transduction

HeLa-JVM and HEK293T cells were plated on a 35 mm dish (Thermo Fisher Scientific, Massachusetts, United States) at 3×10^5^ cells/dish density in DMEM. On the next day (day 0), the cells were transfected with 2 µg DNA (1 µg of shRNA plasmid [pLKO.1-TurboGFP or pALAF034], 0.5 µg of pMD2.G, and 0.5 µg of psPAX2) using 6 µL FuGENE HD (Promega, Wisconsin, United States) in 200 µL Opti-MEM (Gibco) according to the manufacturer’s protocol. On the following day (day 1), the medium was replaced with fresh DMEM. On day 2, the virus-containing medium was collected and filtered through a 0.45 µm filter (Merck Millipore, Massachusetts, United States). The filtered virus medium was used to transduce HeLa-JVM and HEK293T cells.

For virus transduction, cells were plated in a 6-well plate at 2×10^5^ cells/well density in DMEM. The next day (day 0), the medium was replaced with fresh DMEM containing 200 µL of the virus suspension and 8 µg/mL polybrene (Merck Millipore). On day 1, the medium was replaced with fresh DMEM. On day 2, the medium was replaced with fresh DMEM containing 2 µg/mL puromycin (Sigma-Aldrich). The medium was changed daily until all of the uninfected control cells died (typically day 5 post-transduction). The transduced cells were cultured in DMEM containing 1 µg/mL puromycin and used for subsequent analyses.

### Retrotransposition Assays

Retrotransposition assays in cultured cells were performed as described previously with some modifications (19, 47, 52, 74, 76, 77, 81, 84).

For the retrotransposition assay using the pCMV-3Tag-9 overexpression vector in HEK293T cells (Figure 1C), we used an *mEGFPI*-based retrotransposition assay (76). HEK293T cells were plated in 6-well plates (Greiner, Frickenhausen, Germany or Thermo Fisher Scientific) at 2×10^5^ cells/well density in DMEM. On the next day (day 0), the cells were transfected with 1 µg DNA (0.5 µg of cepB-gfp-L1.3, cepB-gfp-L1.3RT[-], or cepB-gfp-L1.3RT[-] intronless and 0.5 µg of pCMV-3Tag-9 control, pALAF023 [HERC5 WT], pALAF062 [HERC5 ΔRLD], pALAF061 [HERC5 ΔHECT], pALAF060 [HERC5 C994A], or pALAF024 [MOV10]; MOV10 served as a positive control) using 3 µL FuGENE HD in 100 µL Opti-MEM according to the manufacturer’s protocol. On the following day (day 1), the medium was replaced with fresh DMEM without antibiotics. On days 2 and 5 post-transfection, the transfected cells were selected with 10 µg/mL Blasticidin S HCl (Sigma-Aldrich). The cells were collected on day 7 post-transfection to determine the percentage of EGFP-positive cells out of 30,000 cells using a flow cytometer, BD FACSCanto II (BD Bioscience, New Jersey, United States) with BD FACSDiva software v.6.1.3 or BD Accuri C6 Plus (BD Bioscience) with FACSAccuri C6 Plus software version 1.0.23.1 (BD Bioscience). The percentage of the resultant EGFP-positive cells was normalized to the transfection efficiency measured by cepB-gfp-L1.3RT(-) intronless EGFP-positive cells, as shown in Figure 1B. Each transfection was carried out with at least two technical replicates.

**Figure 1.**
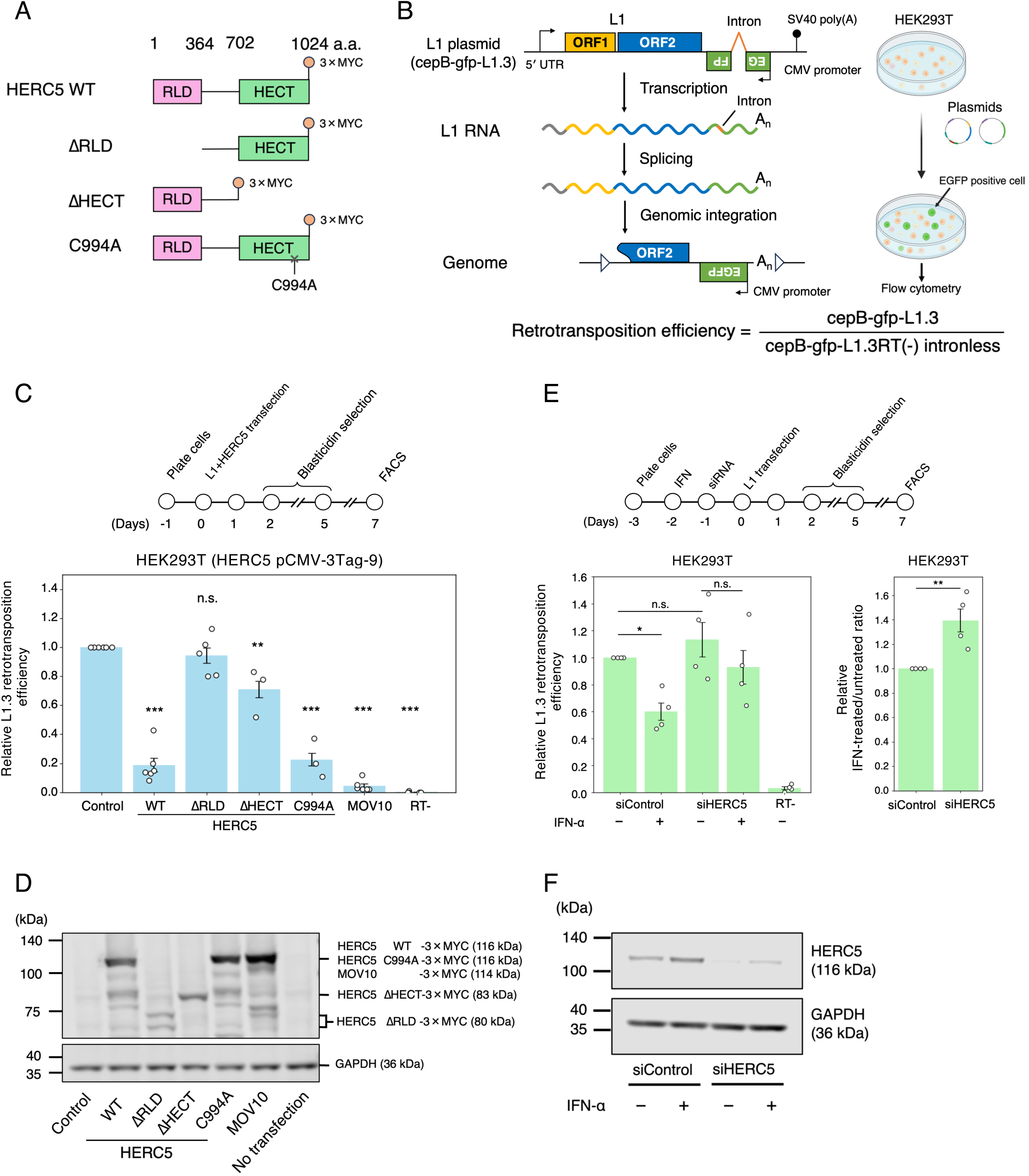
HERC5 inhibits L1 retrotransposition independently of ISGylation. (**A**) Schematic of HERC5 and its mutants used in this study. HERC5 contains an N-terminal RLD domain (pink) and a C-terminal HECT domain (light green). Numbers shown above each domain indicate amino acid (a.a.) positions. A C-terminal 3×MYC epitope tag was fused to HERC5 WT, ΔRLD, ΔHECT, and C994A expression vectors. (**B**) Schematic of the L1 retrotransposition assay. Left: the retrotransposition-competent L1.3-expressing vector (cepB-gfp-L1.3) contains an enhanced green fluorescent protein retrotransposition indicator cassette (*mEGFPI*) in the 3′ UTR of L1 in the opposite orientation relative to the sense strand of L1 transcription. The *mEGFPI* cassette is interrupted by an intron in the same orientation relative to L1 transcription. This ensures that EGFP is expressed only after successful retrotransposition. Right: HEK293T cells were transfected with the L1-expressing plasmid containing the *mEGFPI* indicator cassette. The EGFP-positive cells were quantified using flow cytometry. The figure was created with BioRender. Bottom: retrotransposition efficiency was calculated by normalizing the percentage of EGFP-positive cells obtained in transfection with L1-expressing plasmid (cepB-gfp-L1.3) to that obtained in transfection with an RT-deficient L1-expressing plasmid lacking the intron in *mEGFPI* (cepB-gfp-L1.3RT[-] intronless). (**C**) L1 retrotransposition assay in HEK293T cells using the pCMV-3Tag-9 vector system. Top: timeline of the assay. HEK293T cells were co-transfected with the L1-expressing vector (cepB-gfp-L1.3) and either pCMV-3Tag-9 (control), HERC5 WT, HERC5 mutants, or MOV10 (positive control). Cells independently co-transfected with cepB-gfp-L1.3RT(-) intronless served as transfection-normalization controls. The transfected cells were selected with blasticidin (10 µg/mL), and the percentage of EGFP-positive cells was determined by flow cytometry. Bottom: L1 retrotransposition assay with HERC5 overexpression using pCMV-3Tag-9 in HEK293T. MOV10 and RT-deficient L1 (cepB-gfp-L1.3RT[-]) served as controls. X-axis, name of the transfected constructs. Y-axis, relative L1 retrotransposition efficiency compared to the control (pCMV-3Tag-9, set to 1.0). The error bars represent the mean ± the standard error of the mean (SEM) of at least three independent biological replicates. Each dot represents an independent biological replicate. The *p*-values were calculated using a one-way ANOVA followed by Bonferroni-Holm post-hoc tests; * *p* < 0.05, ** *p* < 0.01, *** *p* < 0.001; n.s.: not significant. (**D**) Protein expression levels of HERC5 WT and mutants in HEK293T. Cells were transfected with HERC5 constructs (pCMV-3Tag-9 vector system). HERC5 and GAPDH proteins were detected by western blot using anti-MYC and anti-GAPDH antibodies, respectively. GAPDH served as a loading control. The predicted molecular weights of the proteins are indicated on the right of the blots. ΔRLD showed two bands, likely due to multiple N-terminal ATG start codons. Although the predicted molecular weight of ΔRLD is ∼80 kDa, it migrated below 75 kDa, probably because of its high glutamic acid and aspartic acid content, consistent with a previous observation (70). (**E**) L1 retrotransposition assay with IFN-α and HERC5 knockdown in HEK293T cells. Top: timeline of the assay. HEK293T cells were treated with IFN-α (100 U/mL) (day −2) and siRNA (day −1) and then transfected with the WT L1-expressing construct (cepB-gfp-L1.3) (day 0). The RT-deficient L1 (cepB-gfp-L1.3RT[-]) served as a negative control. After blasticidin selection (10 µg/mL), the percentage of EGFP-positive cells was measured. Bottom left: L1 retrotransposition assay with IFN-α and siRNA treatments in HEK293T cells. X-axis, siRNA and IFN-α treatments. Y-axis, relative L1 retrotransposition efficiency to the non-targeting siRNA control (siControl) in the absence of IFN-α (set to 1.0). The error bars and *p*-values were calculated as in (C). Each dot represents an independent biological replicate. Bottom right: the relative retrotransposition ratio calculated by normalizing values obtained with IFN-α to those without IFN-α in the left panel. The ratio of siControl is set to 1.0 to compare with that of siHERC5. The *p*-values were calculated using a two-tailed, unpaired Student’s t-test; ** *p* < 0.01. Each dot represents an independent biological replicate. (**F**) Endogenous HERC5 protein expression levels with IFN-α and siRNA treatments in HEK293T cells. HERC5 and GAPDH proteins were detected by western blot using anti-HERC5 and anti-GAPDH antibodies, respectively. GAPDH served as a loading control.

For the retrotransposition assay using the pEBNA overexpression vector in HEK293T cells (Supplementary Figure S1B), we used an *mEGFPI*-based retrotransposition assay (76). HEK293T cells were plated in 6-well plates at 2×10^5^ cells/well density in DMEM. On the next day (day 0), the cells were transfected with 1 µg DNA (0.5 µg of cepB-gfp-L1.3, cepB-gfp-L1.3RT[-], or cepB-gfp-L1.3RT[-] intronless and 0.5 µg of pEBNA control, pKN025 [HERC5 WT], pKN026 [HERC5 ΔRLD], pKN027 [HERC5 ΔHECT], pKN028 [HERC5 C994A], or pALAF024 [MOV10]) using 3 µL FuGENE HD in 100 µL Opti-MEM according to the manufacturer’s protocol. On the following day (day 1), the medium was replaced with fresh DMEM without antibiotics. On days 2 and 5 post-transfection, the transfected cells were selected with 10 µg/mL Blasticidin S HCl. The cells were collected on day 7 post-transfection to determine the percentage of EGFP-positive cells out of 30,000 cells using a flow cytometer, BD Accuri C6 Plus. The percentage of the resultant EGFP-positive cells was normalized to the transfection efficiency measured by cepB-gfp-L1.3RT(-) intronless EGFP-positive cells, as shown in Figure 1B. Each transfection was carried out with at least two technical replicates.

For the retrotransposition assay using HeLa-JVM cells (Supplementary Figure S1C), we used an *mblastI*-based retrotransposition assay (75). HeLa-JVM cells were plated in 6-well plates at 1×10^5^ cells/well density in DMEM. On the next day (day 0), the cells were transfected with 1 µg DNA (0.5 µg of pJJ101 or pJJ105 and 0.5 µg of pCMV-3Tag-9 control, pALAF023 [HERC5 WT], pALAF062 [HERC5 ΔRLD], pALAF061 [HERC5 ΔHECT], pALAF060 [HERC5 C994A], or pALAF024 [MOV10]) using 3 µL FuGENE HD in 100 µL Opti-MEM according to the manufacturer’s protocol. On the following day (day 1), the medium was replaced with fresh DMEM without antibiotics. From day 3 post-transfection, the medium was replaced every 2 days with fresh DMEM containing 10 µg/mL Blasticidin S HCl until no cells remained in the non-transfected control (day 11–13). The Blasticidin-resistant cell foci were washed with cold 1×PBS and fixed with a fixation solution (1×PBS containing 0.2% [v/v] glutaraldehyde [Nacalai Tesque] and 2% [v/v] formaldehyde [Nacalai Tesque]) for 10 min at room temperature. The cells were stained with 0.1% (w/v) crystal violet (Nacalai Tesque), and foci numbers were counted. To measure the transfection efficiency, HeLa-JVM cells were plated in 6-well plates at 2.5×10^3^ cells/well density in DMEM. On the next day (day 0), the cells were transfected with 1 µg DNA (0.5 µg of pcDNA6 and 0.5 µg of pCMV-3Tag-9 control, pALAF023 [HERC5 WT], pALAF062 [HERC5 ΔRLD], pALAF061 [HERC5 ΔHECT], pALAF060 [HERC5 C994A], or pALAF024 [MOV10]) using 3 µL FuGENE HD in 100 µL Opti-MEM according to the manufacturer’s protocol. After ∼24 h, the medium was replaced with fresh DMEM without antibiotics. From day 3 post-DNA transfection, the medium was replaced every 2 days with fresh DMEM containing 10 µg/mL Blasticidin S HCl until day 11 to 13. The blasticidin-resistant cell foci were counted as described above. The L1 retrotransposition efficiency obtained with pJJ101 or pJJ105 was normalized to the transfection efficiency as described in Supplementary Figure S1C. Each transfection was carried out with at least two technical replicates.

For the retrotransposition assay using siRNA with the interferon-α (Abcam, UK, Cambridge) treatment in HEK293T cells (Figure 1E), we used the *mEGFPI*-based retrotransposition assay (76). HEK293T cells were plated in 6-well plates at 2×10^5^ cells/well density in DMEM. The next day (day −2), the cells were treated with 100 U/mL interferon-α or 1×PBS. After ∼24 h (day −1), the medium was replaced with fresh 1 mL DMEM and transfected with 1.25 µL of 20 nM siRNA (Dharmacon, California, United States) (non-targeting control: ON-TARGETplus Non-targeting pool, D-001810-10-0020 or HERC5: ON-TARGETplus HERC5 siRNA SMART pool, L-005174-00-0005) using 3.75 µL of Lipofectamine RNAiMAX (Thermo Fisher Scientific) and 125 µL of Opti-MEM according to the manufacturer’s protocol. On the following day (day 0), the medium was replaced with fresh DMEM, and the cells were transfected with 1 µg DNA (cepB-gfp-L1.3, cepB-gfp-L1.3RT[-] or cepB-gfp-L1.3RT[-] intronless) using 3 µL FuGENE HD in 100 µL Opti-MEM according to the manufacturer’s protocol. The medium was replaced with fresh DMEM without antibiotics on the next day (day 1). On days 2 and 5 post-transfection, DMEM was replaced with fresh DMEM containing 10 µg/mL Blasticidin S HCl. The cells were collected on day 7 post-transfection to determine the percentage of EGFP-positive cells out of 50,000 cells using a flow cytometer, BD Accuri C6 Plus. The percentage of the resultant EGFP-positive cells was normalized to the transfection efficiency measured by cepB-gfp-L1.3RT(-) intronless EGFP-positive cells as described in Figure 1B. Each transfection was carried out with at least two technical replicates.

For the retrotransposition assay using the shRNA-expressing HeLa-JVM cells (see “Virus transduction”, Supplementary Figure S1D), we used an *mEGFPI*-based retrotransposition assay (76). The shRNA-expressing HeLa-JVM cells were plated in 6-well plates at 2×10^5^ cells/well density in DMEM containing 1 µg/mL puromycin. On the next day (day 0), the cells were transfected with 0.5 µg DNA (cepB-gfp-L1.3, cepB-gfp-L1.3RT[-] or cepB-gfp-L1.3RT[-] intronless) using 1.5 µL FuGENE HD in 100 µL Opti-MEM according to the manufacturer’s protocol. On the following day (day 1), the medium was replaced with fresh DMEM containing 1 µg/mL puromycin. On days 2 and 5 post-transfection, DMEM was replaced with fresh DMEM containing 10 µg/mL Blasticidin S HCl and 1 µg/mL puromycin. The cells were collected on day 7 post-transfection to determine the percentage of EGFP-positive cells out of 30,000 cells using a flow cytometer, BD FACSCalibur (BD Bioscience) with software CellQuest Pro v5.2 (BD Bioscience). The percentage of the resultant EGFP-positive cells was normalized to the transfection efficiency measured by cepB-gfp-L1.3RT(-) intronless EGFP-positive cells as described in Figure 1B. Each transfection was carried out with at least two technical replicates.

For the *Alu* and L1 retrotransposition assay using HeLa-HA cells (Figure 4D, E, and Supplementary Figure S4B), we used an *mneoI*-based retrotransposition assay (19, 52). HeLa-HA cells were plated in 6-well plates at 2×10^5^ cells/well density in MEM medium. On the next day (day 0), the cells were transfected with 1 µg DNA (for *Alu* retrotransposition assay; Figure 4D: 0.5 µg of pKN040, pKN041, or phrGFP-C and 0.5 µg of pCMV-3Tag-8-Barr, pALAF023 [HERC5], or pALAF024 [MOV10], Figure 4E: 0.5 µg of pTM489, pKO001, or phrGFP-C and 0.5 µg of pCMV-3Tag-8-Barr, pALAF023 [HERC5], or pALAF024 [MOV10], for L1 retrotransposition assay; Supplementary Figure S4B: 0.5 µg of pJM101/L1.3, pJM105/L1.3, or phrGFP-C and 0.5 µg of pCMV-3Tag-8-Barr, pALAF023 [HERC5], or pALAF024 [MOV10]) using 3 µL FuGENE HD in 100 µL Opti-MEM according to the manufacturer’s protocol. On the following two days (days 1 and 2), the medium was replaced with fresh MEM without antibiotics. From day 3 onwards, the medium was replaced daily with fresh MEM containing 500 µg/mL G418 until no cells remained in the non-transfected control (day 15). The G418-resistant cell foci were washed with cold 1×PBS, and fixed with the fixation solution for 10 min at room temperature. The cells were stained with 0.1% (w/v) crystal violet, and foci numbers were counted. To measure the transfection efficiency, HeLa-HA cells were plated in 6-well plates at 2×10^5^ cells/well density in MEM medium. On the next day (day 0), the cells were transfected with 1 µg DNA (0.5 µg of phrGFP-C and 0.5 µg of pCMV-3Tag-8-Barr, pALAF023 [HERC5], or pALAF024 [MOV10]) using 3 µL FuGENE HD in 100 µL Opti-MEM according to manufacturer’s protocol. On day 3 post-transfection, cells were washed with cold 1×PBS, trypsinized with 0.25% (v/v) Trypsin-EDTA (Gibco), and harvested to count EGFP-positive cell numbers out of 30,000 cells using the flow cytometer, BD FACSCalibur. The percentage of G418-resistant foci was normalized to the transfection efficiency measured by phrGFP-C EGFP-positive cells, instead of the pcDNA6 transfection efficiency. Each transfection was carried out with at least two technical replicates.

For the retrotransposition assay of mouse L1 species in HEK293T cells (Figure 6A and B), we used an *mEGFPI*-based retrotransposition assay (76). HEK293T cells were plated in 6-well plates at 2×10^5^ cells/well density in DMEM high glucose medium. On the next day (day 0), the cells were transfected with 1 µg DNA (Figure 6A: 0.5 µg of cep99-gfp-TG_F_21 [TG_F_21 WT], cep99-gfp-L1.3RT[-] [L1.3 RT−], or cep99-gfp-L1.3RT[-] intronless and 0.5 µg of pCMV-3Tag-8-Barr, pALAF023 [HERC5], or pALAF024 [MOV10], Figure 6B: 0.5 µg of cep99-gfp-ORFeus-Mm [ORFeus-Mm WT], cep99-gfp-ORFeus-Mm mut2 [EN− RT−], or cep99-gfp-L1.3RT[-] intronless and 0.5 µg of pCMV-3Tag-8-Barr, pALAF023 [HERC5], or pALAF024 [MOV10]) using 3 µL FuGENE HD in 100 µL Opti-MEM according to the manufacturer’s protocol. On the following day (day 1), the medium was replaced with fresh DMEM high glucose medium without antibiotics. On days 2 and 5 post-transfection, the transfected cells were selected with 1 µg/mL puromycin. The cells were collected on day 7 post-transfection to determine the percentage of EGFP-positive cells out of 30,000 cells using a flow cytometer, BD FACSCalibur. The percentage of the resultant EGFP-positive cells was normalized to the transfection efficiency measured by cep99-gfp-L1.3RT(-) intronless EGFP-positive cells as described in Figure 1B. Each transfection was carried out with at least two technical replicates.

For the retrotransposition assay of zebrafish L2 in HEK293T cells (Figure 6C), we used an *mEGFPI*-based retrotransposition assay (76). HEK293T cells were plated in 6-well plates at 5.0×10^5^ cells/well density in DMEM high glucose medium. On the next day (day 0), the cells were transfected with 1 µg DNA (0.5 µg of cep99-gfp-Z2 [ZfL2-2 WT], cep99-gfp-Z2-RTm [ZfL2-2 RT−], or cep99-gfp-L1.3RT[-] intronless and 0.5 µg of pCMV-3Tag-8-Barr, pALAF023 [HERC5], or pALAF024 [MOV10]) using 3 µL FuGENE HD in 100 µL Opti-MEM according to the manufacturer’s protocol. On the following day (day 1), the medium was replaced with fresh DMEM high glucose medium without antibiotics. From day 2 onwards, the medium was replaced with fresh DMEM high glucose medium containing 1 µg/mL puromycin until day 5. The cells were collected on day 5 post-transfection to determine the percentage of EGFP-positive cells out of 30,000 cells using a flow cytometer, BD FACSCalibur. The percentage of the resultant EGFP-positive cells was normalized to the transfection efficiency measured by cep99-gfp-L1.3RT(-) intronless EGFP-positive cells, as shown in Figure 1B. Each transfection was carried out with at least two technical replicates.

For the retrotransposition assay of L1 ORFeus (Figure 3D), HEK293T cells were plated in 6-well plates at 2×10^5^ cells/well density in DMEM. On the next day (day 0), the cells were transfected with 1 µg DNA (0.5 µg of pKN036 [cepB-gfp-L1 ORFeus], cepB-gfp-L1.3, cepB-gfp-L1.3RT[-], or cepB-gfp-L1.3RT[-] intronless and 0.5 µg of pCMV-3Tag-9 or pALAF023 [HERC5]) using 3 µL FuGENE HD in 100 µL Opti-MEM according to the manufacturer’s protocol. On the following day (day 1), the medium was replaced with fresh DMEM without antibiotics. On days 2 and 5 post-transfection, the transfected cells were selected with 10 µg/mL Blasticidin S HCl. The cells were collected on day 7 post-transfection to determine the percentage of EGFP-positive cells out of 30,000 cells using a flow cytometer, BD Accuri C6 Plus. The percentage of the resultant EGFP-positive cells was normalized to the transfection efficiency measured by cepB-gfp-L1.3RT(-) intronless EGFP-positive cells, as shown in Figure 1B. Each transfection was carried out with at least two technical replicates.

For the retrotransposition assay of the small HERC family in HEK293T cells (Figure 6F), HEK293T cells were plated in 6-well plates at 2×10^5^ cells/well density in DMEM. On the next day (day 0), the cells were transfected with 1 µg DNA (0.5 µg of cepB-gfp-L1.3, cepB-gfp-L1.3RT[-], or cepB-gfp-L1.3RT[-] intronless and 0.5 µg of pCMV-3Tag-9, pKN031 [HERC4], pALAF023 [HERC5], pKN032 [hHERC6], or pKN036 [mHERC6]) using 3 µL FuGENE HD in 100 µL Opti-MEM according to the manufacturer’s protocol. On the following day (day 1), the medium was replaced with fresh DMEM without antibiotics. On days 2 and 5 post-transfection, the transfected cells were selected with 10 µg/mL Blasticidin S HCl. The cells were collected on day 7 post-transfection to determine the percentage of EGFP-positive cells out of 30,000 cells using a flow cytometer, BD Accuri C6 Plus. The percentage of the resultant EGFP-positive cells was normalized to the transfection efficiency measured by cepB-gfp-L1.3RT(-) intronless EGFP-positive cells, as shown in Figure 1B. Each transfection was carried out with at least two technical replicates.

### RNA extraction and RT-qPCR

For the quantification of L1 RNA (Figure 2B), HEK293T cells were plated into two wells of 6-well plates at 2×10^5^ cells/well density in DMEM. On the next day (day 0), the cells were transfected with 1.5 µg of DNA (0.5 µg of L1-expressing plasmid [pTMF3] and 1 µg of pCMV-3Tag-9, pALAF023 [HERC5], or pALAF015 [HELZ2]) using 3 µL FuGENE HD in 100 µL Opti-MEM according to the manufacturer’s protocol. The following day (day 1), the medium was replaced with fresh DMEM without antibiotics. On day 3 post-transfection, HEK293T cells were harvested by pipetting and washed twice with cold 1×PBS. The cell pellets were flash frozen with liquid nitrogen and stored at −80°C for subsequent experiments. The RNA was extracted with the QIAshredder (QIAGEN, Venlo, Netherlands) and RNeasy Plus Mini Kit (QIAGEN) according to the manufacturer’s protocol. Total RNA was eluted in 30 µL of RNase-free water, and the RNA concentration was measured.

**Figure 2.**
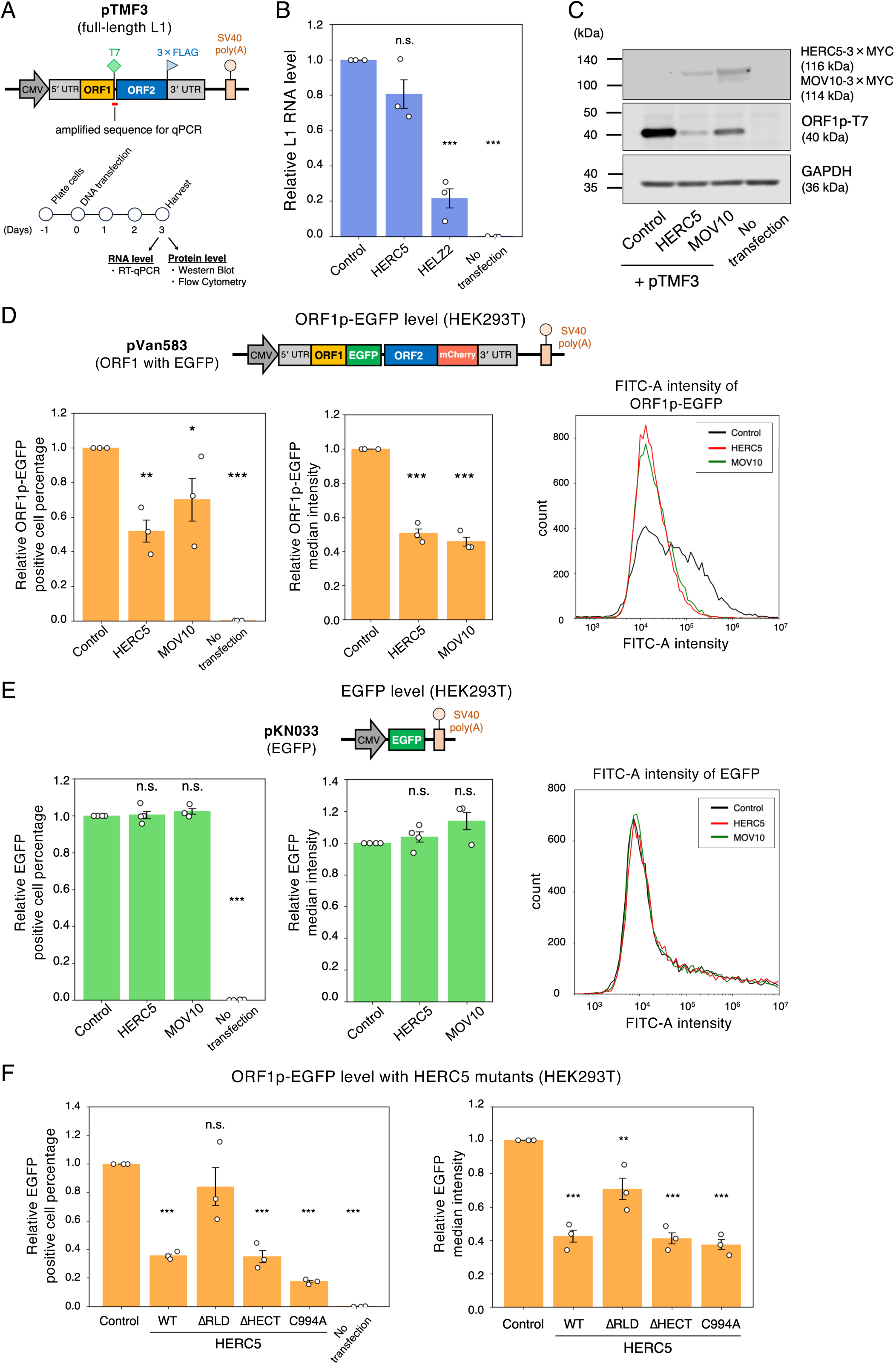
HERC5 reduces L1 ORF1p without affecting L1 RNA levels. (**A**) Experimental design for checking L1 RNA and ORF1p levels. Top: schematic of the full-length L1 construct (pTMF3), which expresses ORF1p tagged with a T7 gene 10 epitope and ORF2p tagged with a 3×FLAG epitope at their carboxyl termini. The red bar indicates the amplified region for the RT-qPCR primer pair used to measure L1 RNA levels (primer sequences are described in Materials and Methods). Bottom: timeline of the experiment. HEK293T cells were co-transfected with L1 expression constructs and either pCMV-3Tag-9 (control), HERC5, HELZ2 (positive control for RNA levels) or MOV10 (positive control for protein levels). The cells were harvested 3 days post-transfection. L1 RNA levels were measured by RT-qPCR, and ORF1p levels were measured by western blot or flow cytometry. (**B**) L1 RNA levels with HERC5. The T7 primer pair was used to quantify L1 RNA levels, which were normalized to GAPDH RNA levels. X-axis, name of the transfected constructs. Y-axis, relative L1 RNA level compared to the control (pCMV-3Tag-9, set to 1.0). The error bars represent the mean ± the standard error of the mean (SEM) of three independent biological replicates. Each dot represents an independent biological replicate. The *p*-values were calculated using a one-way ANOVA followed by Bonferroni-Holm post-hoc tests; * *p* < 0.05, ** *p* < 0.01, *** *p* < 0.001; n.s.: not significant. (**C**) ORF1p levels with HERC5. HERC5 and MOV10 were detected by an anti-MYC antibody. ORF1p and GAPDH were detected by anti-T7 and anti-GAPDH antibodies, respectively. GAPDH served as a loading control. (**D**) Flow cytometry analysis of ORF1p-EGFP expression. HEK293T cells were co-transfected with pVan583 and either pCMV-3Tag-9, HERC5, or MOV10 expression vector. Top: schematic of the pVan583 plasmid, which expresses EGFP-tagged ORF1p and mCherry-tagged ORF2p and contains the CMV promoter, the 5′ UTR promoter, and the SV40 poly(A) signal sequence for L1 expression. Bottom left and middle panels: relative percentages and median intensities (FITC-A) of ORF1p-EGFP-positive cells, respectively. X-axis, name of the transfected constructs. Y-axis, relative percentages or median intensities compared to the control (pCMV-3Tag-9, set to 1.0). The error bars and *p*-values were calculated as in (B). Bottom right: overlaid frequency distribution plot of FITC-A intensity of control (black), HERC5 (red), or MOV10 (green). X-axis, log-scaled fluorescence intensity. Y-axis, cell count. (**E**) Flow cytometry analysis measuring EGFP expression. HEK293T cells were co-transfected with pKN033 and either pCMV-3Tag-9, HERC5, or MOV10. Top: schematic of the pKN033 plasmid, which is derived from pVan583 and expresses EGFP alone. Bottom left and middle panels: relative percentages and median intensities of EGFP-positive cells, respectively. X-axis, name of the transfected constructs. Y-axis, relative percentages or median intensities compared to the control (pCMV-3Tag-9, set to 1.0). The error bars and *p*-values were calculated as in (B). Bottom right: overlaid frequency distribution plot of FITC-A intensity of control (black), HERC5 (red), or MOV10 (green). X-axis, log-scaled fluorescence intensity. Y-axis, cell count. (**F**) ORF1p-EGFP levels with HERC5 WT and its mutants. HEK293T cells were co-transfected with pVan583 and either pEBNA, HERC5 WT, or its mutants. The relative cell percentages and median intensities of ORF1p-EGFP-positive cells are shown in the left and middle panels, respectively. X-axis, name of the transfected constructs. Y-axis, relative percentages or median intensities compared to the control (pEBNA, set to 1.0). The error bars and *p*-values were calculated as in (B).

One microgram of total RNA was used as a template for the following reverse transcription reaction using 0.2 mM dNTP (Takara Bio, Shiga, Japan), 1 U/µL ribonuclease inhibitor (Takara Bio), 0.25 U/µL AMV reverse transcriptase XL (Takara Bio), and 0.125 µM Oligo (dT) primer (Eurofins, Luxembourg). Two negative controls were included: RT−, in which reverse transcriptase was omitted from the reaction, and NT− (no template), in which template RNA was replaced with RNase-free water for reverse transcription. The reverse transcription reaction was performed as follows: 30°C for 10 min, 42°C for 30 min, 95°C for 5 min.

qPCR was performed using Luna Universal qPCR Master Mix (New England Biolabs, Massachusetts, United States) and HT7900 Fast Real-time PCR System (Applied Biosystems, Massachusetts, United States). Real-time qPCR was performed as follows: 95°C for 1 min; then 40 cycles of 95°C for 15 s (denaturation), and 60°C for 1 min (amplification). Technical duplicates were run for each sample. The quantification of each cDNA was determined by comparing the cycle threshold (Ct) value with a standard curve generated from one of the samples using the Sequence Detection Systems software version 2.4.2 (Applied Biosystems). The Ct values were all detected within the range of the standard curves used in this study.

### Primers used for RT-qPCR

L1 (SV40)_F: 5′-TCCAGACATGATAAGATACATTGATGAG-3′

L1 (SV40)_R: 5′-GCAATAGCATCACAAATTTCACAAA-3′

L1 (T7)_F: 5′-ATGGCTAGCATGACTGGTGG-3′

L1 (T7)_R: 5′-CCTGTCATTATGATGTTAGCTGGTG-3′

GAPDH_F: 5′-GGAGTCCCTGCCACACTCAG-3′

GAPDH_R: 5′-GGTCTACATGGCAACTGTGAGG-3′

Oligo (dT): 5′-TTTTTTTTTTTTTTTTTTTTVN-3′

### Western blots

For the detection of overexpressed protein levels (Figure 2C, 3E, 4A, B, C, Supplementary Figure S1B, S2A, S3C, and S4A), HEK293T cells were plated into two wells of 6-well plates at 2×10^5^ cells/well density in DMEM. On the next day (day 0), the cells were transfected with DNA (L1-expressing plasmid and/or ISG-expressing plasmids; Figure 1D: 1 µg of pCMV-3Tag-9, pALAF023 [HERC5 WT], pALAF062 [HERC5 ΔRLD], pALAF061 [HERC5 ΔHECT], pALAF060 [HERC5 C994A], Figure 2C: 0.5 µg of L1-expressing plasmid [pTMF3] and 1 µg of pCMV-3Tag-9, pALAF023 [HERC5], or pALAF024 [MOV10], Figure 3E: 0.5 µg of L1-expressing plasmid [pJM101/L1.3 or pKN024] and 1 µg of pCMV-3Tag-9 or pALAF023 [HERC5], Figure 4A: 0.5 µg of L1-expressing plasmid [pKN035] and 1 µg of pCMV-3Tag-9, pALAF023 [HERC5], or pALAF024 [MOV10], Figure 4B: 0.5 µg of L1-expressing plasmid [pTMO2F3] and 1 µg of pCMV-3Tag-9 or pALAF023 [HERC5], Figure 4C: 0.5 µg of L1-expressing plasmid [pTMF3] and 1 µg of pCMV-3Tag-9 or pALAF023 [HERC5], Supplementary Figure S1B: pEBNA, pKN025 [HERC5 WT], pKN026 [HERC5 ΔRLD], pKN027 [HERC5 ΔHECT], pKN028 [HERC5 C994A], or pALAF024 [MOV10], Supplementary Figure S2A: 1 µg of L1-expressing plasmid [pJM101/L1.3], Supplementary Figure S3C: 0.5 µg of L1-expressing plasmid [pJM101/L1.3FLAG or pALAF008] and 1 µg of pCMV-3Tag-9 or pALAF023 [HERC5], and Supplementary Figure S4A: 0.5 µg of L1-expressing plasmid [pTMF3, pTMF3_Δ5UTR, or pL1(5&3UTRs_Fluc)] and 1 µg of pCMV-3Tag-9 or pALAF023 [HERC5]) using 3 µL FuGENE HD in 100 µL Opti-MEM according to the manufacturer’s protocol. On the following day (day 1), the medium was replaced with fresh DMEM. On day 3 post-transfection, HEK293T cells were harvested by pipetting and washed twice with cold 1×PBS. The cell pellets were flash frozen with liquid nitrogen and stored at −80°C for subsequent experiments.

**Figure 3.**
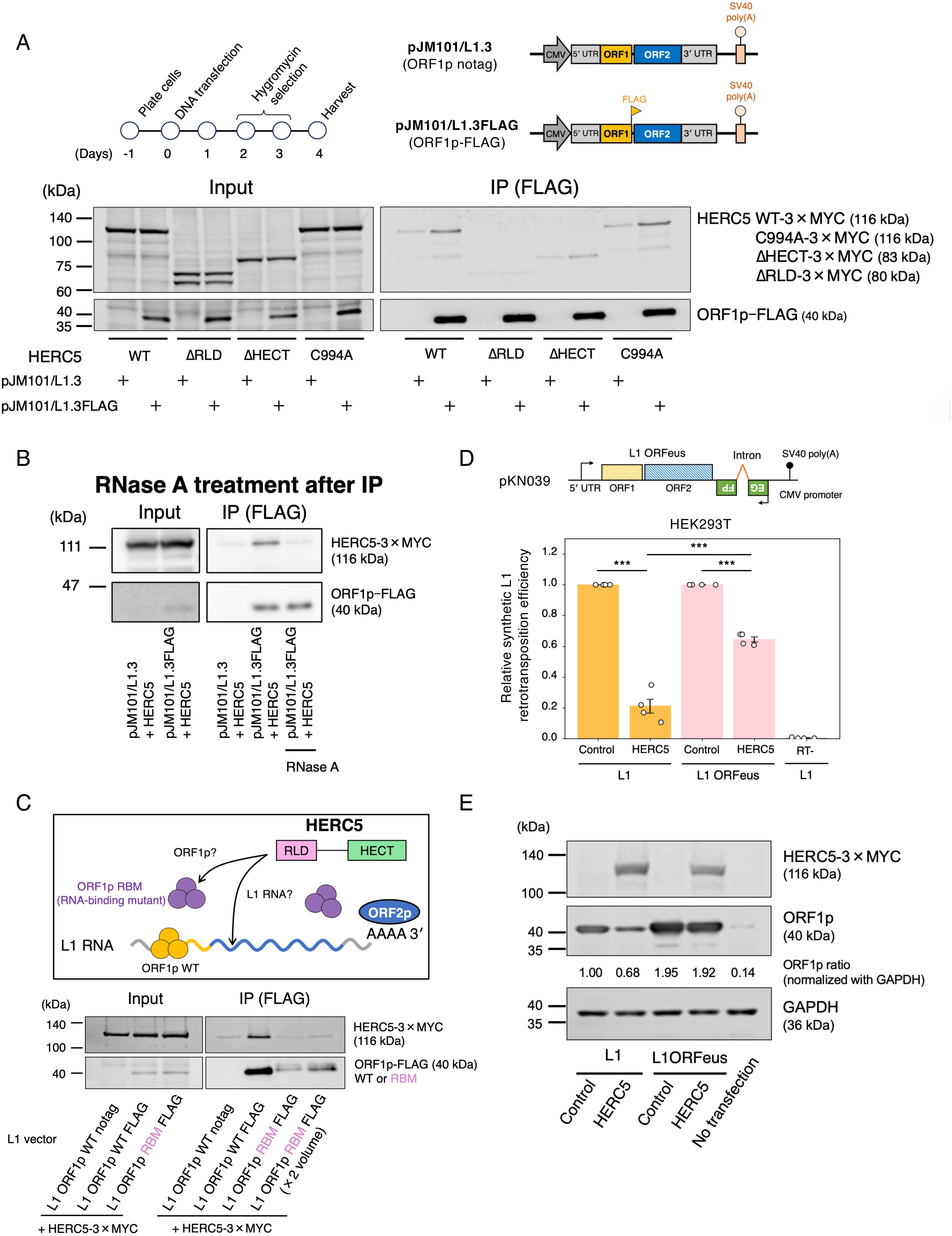
HERC5 interacts with L1 RNA. (**A**) Interaction of ORF1p with HERC5 WT and its mutants. Top left: timeline of the experiment. HEK293T cells were co-transfected with L1 and HERC5 expression vectors. Cells were harvested on 4 days post-transfection, and ORF1p-FLAG complexes were immunoprecipitated. Top right: schematic of L1-expressing plasmids. The pJM101/L1.3 plasmid contains the full-length L1.3, and pJM101/L1.3FLAG expresses ORF1p tagged with a FLAG epitope at the carboxyl terminus. Bottom: the input and anti-FLAG IP reactions were analyzed by western blotting. pJM101/L1.3 served as a negative control. HERC5 and its mutants expressed from pEBNA were detected by an anti-MYC antibody, and ORF1p was detected by an anti-FLAG antibody. (**B**) Co-immunoprecipitation of ORF1p and HERC5 with RNase treatment. HEK293T cells were co-transfected with L1 and HERC5 expression vectors. ORF1p-FLAG complexes were purified as in (A) and were treated in the presence or absence of RNase A. The rightmost lane shows the RNase A-treated ORF1p-FLAG complex. HERC5 and ORF1p were detected by anti-MYC and anti-FLAG antibodies, respectively. (**C**) Interaction of the ORF1p RNA-binding mutant (RBM) with HERC5. HEK293T cells were co-transfected with HERC5 and either L1 WT or RBM expression vectors. Top: schematic diagram of L1 RNP with ORF1p WT (yellow) and RBM (purple). Bottom: the input and anti-FLAG IP reactions were analyzed by western blotting. FLAG-tagged ORF1p WT and RBM were immunoprecipitated. HERC5 and ORF1p were detected by anti-MYC and anti-FLAG antibodies, respectively. (**D**) Retrotransposition assay of L1 ORFeus and L1.3. Top: schematic of the pKN039 plasmid, which expresses codon-optimized ORF1p and ORF2p. L1 ORFeus is driven by the CMV promoter and the L1.3 5′ UTR promoter and terminates with the SV40 poly(A) signal sequence. The *mEGFPI* retrotransposition indicator cassette was inserted into the L1 3′ UTR. Bottom: the RT-deficient L1 (cepB-gfp-L1.3RT[-]) served as a negative control. X-axis, name of the transfected constructs. Y-axis, relative retrotransposition efficiency compared to the control (pCMV-3Tag-9 was set to 1.0 for each L1 construct). The error bars represent the mean ± the standard error of the mean (SEM) of four independent biological replicates. Each dot represents an independent biological replicate. The *p*-values were calculated using a one-way ANOVA followed by Bonferroni-Holm post-hoc tests; *** *p* < 0.001; n.s.: not significant. (**E**) ORF1p levels from L1.3 and L1 ORFeus with HERC5. HEK293T cells were transfected with L1 ORFeus and either the empty vector (pCMV-3Tag-9) or a HERC5-expressing vector. HERC5, ORF1p, and GAPDH were detected by anti-MYC, anti-ORF1p, and anti-GAPDH antibodies, respectively. ORF1p and GAPDH band signal intensities were measured with Empiria Studio. ORF1p signal intensities were normalized to GAPDH intensities to calculate the ORF1p ratio. The signal intensity of L1.3 ORF1p in the control condition was set to 1.0. GAPDH served as a loading control.

**Figure 4.**
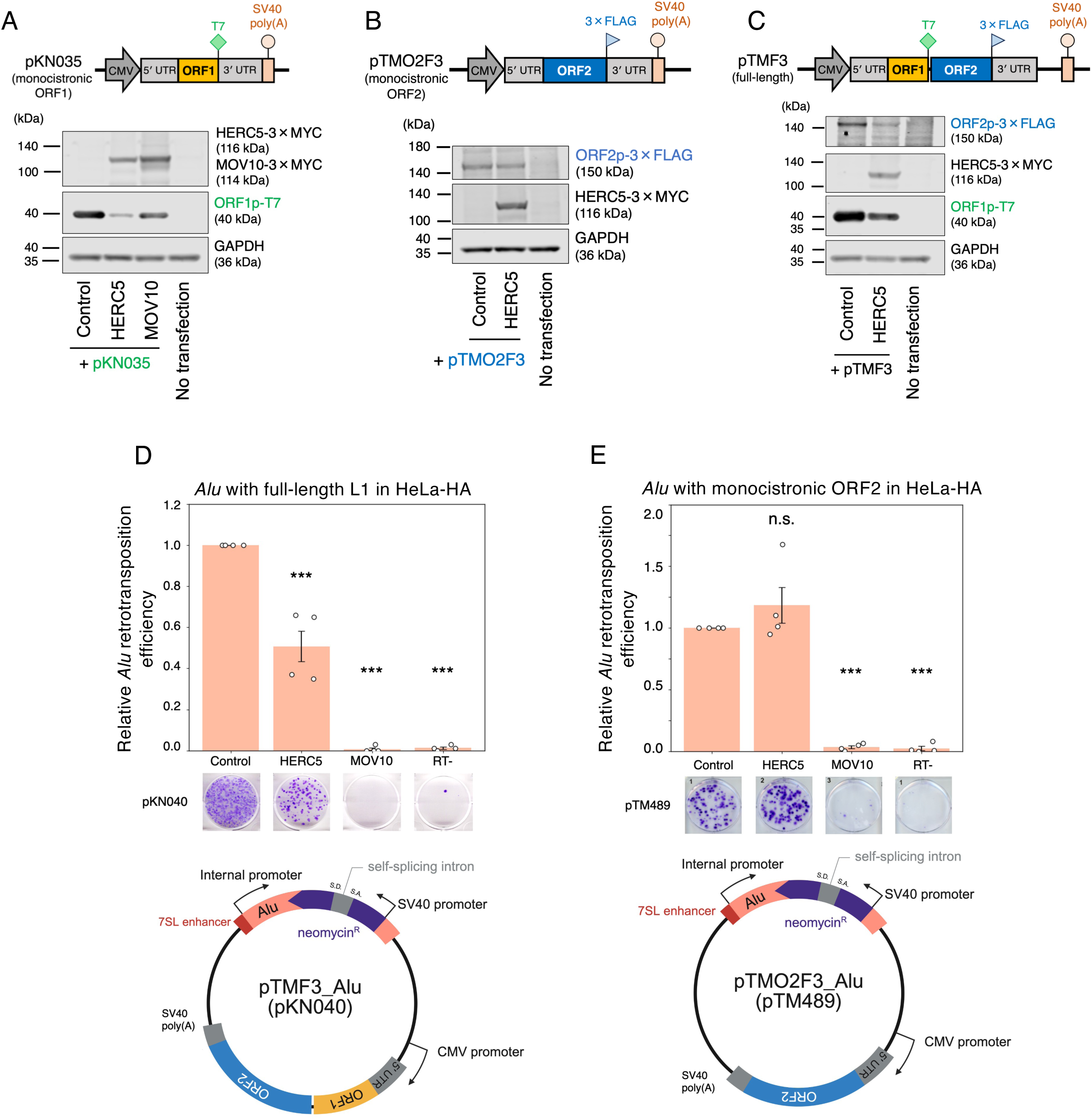
HERC5 targets ORF1p and its downstream ORF2p. (**A**–**C**) Protein levels of modified L1 constructs with HERC5. HEK293T cells were co-transfected with modified L1 and HERC5-expressing vectors. The cells were harvested 3 days post-transfection, and protein levels were assessed by western blotting. Top: schematic of modified L1 vectors. Left: schematic of pKN035 (A), which expresses monocistronic ORF1p tagged with a T7 gene 10 epitope. Middle: schematic of pTMO2F3 (B), which expresses monocistronic ORF2p with a 3×FLAG epitope. Right: schematic of pTMF3 (C), which expresses ORF1p tagged with a T7 gene 10 epitope and ORF2p tagged with a 3×FLAG epitope at their carboxyl termini. Bottom: the protein expression levels were assessed by western blotting. HERC5 and MOV10 were detected by an anti-MYC antibody. ORF1p, ORF2p, and GAPDH were detected by anti-T7, anti-FLAG, and anti-GAPDH antibodies, respectively. GAPDH served as a loading control. (**D**) *Alu* retrotransposition assay with full-length L1 in HeLa-HA. Top: HeLa-HA cells were co-transfected with the *Alu* and full-length L1-expressing construct (pKN040) and either pCMV-3Tag-8-Barr (control), a HERC5-expressing construct, or a MOV10-expressing construct. Cells were selected with G418 (500 µg/mL), stained with crystal violet, and the resulting colonies were counted. The representative images of stained G418-resistant colonies are shown below each condition. The colony numbers of pKN040 were normalized to transfection efficiency and determined as retrotransposition efficiency. MOV10 and the RT mutant served as controls. X-axis, name of the transfected constructs. Y-axis, relative Alu retrotransposition efficiency compared to the control (pCMV-3Tag-8-Barr, set to 1.0). The error bars represent the mean ± the standard error of the mean (SEM) of four independent biological replicates. Each dot represents an independent biological replicate. The *p*-values were calculated using a one-way ANOVA followed by Bonferroni-Holm post-hoc tests; *** *p* < 0.001; n.s.: not significant. Bottom: schematic of pKN040. The *Alu* sequence contains the neomycin-resistant gene cassette in the opposite direction of *Alu* transcription, and the self-splicing intron was inserted into the neomycin-resistant gene in the same direction as the *Alu* sequence, together with the full-length L1 sequence. S.D.: splice donor site, S.A.: splice acceptor site. The plasmid figure was created with BioRender. (**E**) *Alu* retrotransposition assay with monocistronic ORF2 in HeLa-HA. Top: HeLa-HA cells were co-transfected with the *Alu* and monocistronic ORF2-expressing construct (pTM489) and either pCMV-3Tag-8-Barr, a HERC5-expressing construct, or a MOV10-expressing construct. The assay was conducted as noted in (D). X-axis, name of the transfected constructs. Y-axis, relative retrotransposition efficiency compared to the control (pCMV-3Tag-8-Barr, set to 1.0). MOV10 and the EN/RT mutant served as controls. The error bars and *p*-values were calculated as in (D). Bottom: schematic of pTM489, which is similar to pKN040, but does not contain the *ORF1* sequence.

For the detection of endogenous HERC5 protein levels with siRNA knockdown and IFN-α treatment (Figure 1F), HEK293T cells were plated into two wells of 6-well plates at 2×10^5^ cells/well density in DMEM (day −2). The next day (day −1), the cells were treated with 100 U/mL IFN-α or 1×PBS. After ∼24 h (day 0), the medium was replaced with fresh 1 mL DMEM and transfected with 1.25 µL of 20 nM siRNA using 3.75 µL of Lipofectamine RNAiMAX and 125 µL of Opti-MEM according to the manufacturer’s protocol. On the following day (day 1), the medium was replaced with fresh DMEM. On day 2 post-transfection, HEK293T cells were harvested by pipetting and washed twice with cold 1×PBS. The cell pellets were flash frozen with liquid nitrogen and stored at −80°C for subsequent experiments.

For the detection of endogenous HERC5 protein levels in shRNA knockdown cells (Supplementary Figure S1E), the control (HeLa-JVM shControl or HEK293T shControl) and HERC5 knockdown (HeLa-JVM shHERC5 or HEK293T shHERC5) cells were plated into two wells of 6-well plates at 2×10^5^ cells/well density in DMEM containing 1 µg/mL puromycin (day 0). On day 2, the medium was replaced with fresh DMEM containing 1 µg/mL puromycin. On the next day (day 3), the HeLa-JVM cells were washed with cold 1×PBS and trypsinized with 0.25% Trypsin-EDTA. HEK293T cells were harvested by pipetting and washed twice with cold 1×PBS. The cell pellets were flash frozen with liquid nitrogen and stored at −80°C for subsequent experiments.

The cells were lysed with Radio-Immunoprecipitation Assay (RIPA) buffer (10 mM Tris-HCl [pH 7.5] [Nacalai Tesque], 1 mM EDTA [Nacalai Tesque], 1% [v/v] Triton X-100 [Nacalai Tesque], 0.1% [w/v] sodium deoxycholate [Nacalai Tesque], 0.1% [w/v] SDS [Nacalai Tesque], and 140 mM NaCl [Nacalai Tesque]) containing 1×cOmplete EDTA-free protease inhibitor cocktail (Roche Diagnostics, Basel, Switzerland) on ice for 30 min. The insoluble cell debris was pelleted at 12,000 rpm for 5 min at 4°C using MDX-310 (Tomy Seiko, Tokyo, Japan). The protein concentration of all resultant supernatants was measured using Bradford Protein Assay (Bio-Rad, California, United States) and subsequently adjusted to the same level. The protein lysate was mixed with an equal volume of 3×SDS sample buffer (187.5 mM Tris-HCl [pH 6.8], 30% [v/v] glycerol [Nacalai Tesque], 6% [w/v] SDS, 0.3 M DTT [Nacalai Tesque], 0.02% (w/v) bromophenol blue [Nacalai Tesque]) and boiled at 100°C for 5 min. Equal amounts of total protein were loaded onto 5–20% Extra PAGE One Precast Gel (Nacalai Tesque) and separated by sodium dodecyl sulfate-polyacrylamide gel electrophoresis (SDS-PAGE). For wet transfer, proteins in the gels were transferred onto 0.45 µm pore Immobilon-FL polyvinylidene difluoride (PVDF) transfer membranes (Merck Millipore) using 10 mM CAPS buffer (3-[cyclohexylamino] propane-1-sulfonic acid [Nacalai Tesque], pH 11.0) in a Mini Trans-Blot Electrophoretic Transfer Cell tank (Bio-Rad) at 4°C at 50 V for 14 h. For semi-dry transfer, proteins were transferred onto 0.45 µm pore Immobilon-FL PVDF transfer membranes using semi-dry buffer (24 mM Tris-HCl, 0.1% [w/v] SDS, 192 mM glycine [Nacalai Tesque], 20% [v/v] ethanol [Nacalai Tesque]) in Trans-Blot SD Semi-Dry Transfer Cell (Bio-Rad) at room temperature, at 10 V for 1 h. After the transfer, the membranes were incubated with 1×Tris-NaCl-Tween (TNT) buffer (0.1 M Tris-HCl [pH 7.5], 140 mM NaCl, 0.1% [v/v] Tween 20 [Nacalai Tesque]) containing 3% [w/v] skim milk (MEGMILK SNOW BRAND, Tokyo, Japan) for 30 min. After washing the membranes with 1×TNT buffer four times for 5 min each, the membranes were incubated with relevant primary antibodies at 4°C overnight. The next day, the membranes were washed with 1×TNT buffer four times for 5 min each and incubated with relevant secondary antibodies in 1×TNT buffer containing 0.01% (w/v) SDS at room temperature for 1 h. The membranes were washed with 1×TNT buffer four times for 5 min per wash. Signals were detected with the Odyssey DLx Imaging System (LI-COR, Nebraska, United States) and analyzed with Empiria Studio Software version 3.2.0.186 (LI-COR).

### EGFP-tagged ORF1p measurement by flow cytometry

For the detection of EGFP-tagged ORF1p-positive cell percentage and its intensity by flow cytometry (Figure 2D, E, and F), HEK293T cells were plated into 6-well plates at 2×10^5^ cells/well density in DMEM. On the next day (day 0), the cells were transfected with 1.5 µg DNA (0.5 µg of an EGFP-tagged ORF1p-expressing plasmid or an EGFP alone-expressing plasmid and 1 µg of HERC5-expressing plasmids; Figure 2D: 0.5 µg of EGFP-tagged ORF1p-expressing plasmid [pVan583] and 1 µg of pCMV-3Tag-9, pALAF023 [HERC5], or pALAF024 [MOV10], Figure 2E: 0.5 µg of the EGFP alone-expressing plasmid [pKN033] and 1 µg of pCMV-3Tag-9, pALAF023 [HERC5], or pALAF024 [MOV10], Figure 2F: 0.5 µg of pVan583 and 1 µg of pEBNA, pKN025 [HERC5 WT], pKN026 [HERC5 ΔRLD], pKN027 [HERC5 ΔHECT], or pKN028 [HERC5 C994A]) using 3 µL FuGENE HD in 100 µL Opti-MEM according to the manufacturer’s protocol. On the following day (day 1), the medium was replaced with fresh DMEM. On day 3 post-transfection, HEK293T cells were harvested by pipetting and washed twice with cold 1×PBS. The percentage of EGFP-positive cells was measured from 30,000 cells, and the median intensity was measured from 10,000 EGFP-positive cells using a flow cytometer, BD Accuri C6 Plus.

### Immunoprecipitation for western blotting

For the immunoprecipitation coupled with western blotting to check the interaction of ORF1p and HERC5 mutants (Figure 3A), HEK293T cells were plated in 10 cm dishes (Thermo Fisher Scientific) at 3×10^6^ cells/dish density in DMEM. On the next day (day 0), the cells were transfected with 6 µg DNA (4 µg of pJM101/L1.3 or pJM101/L1.3FLAG and 2 µg of pKN025 [HERC5 WT], pKN026 [HERC5 ΔRLD], pKN027 [HERC5 ΔHECT], or pKN028 [HERC5 C994A]) using 18 µL of 1 mg/mL transfection-grade linear poly-ethylenimine hydrochloride (MW 40,000) (PEI-MAX-40K) (Polysciences, Warrington, United States) in 500 µL Opti-MEM. On the following day (day 1), the medium was replaced with fresh DMEM. On days 2 and 3 post-transfection, the medium was replaced with fresh DMEM containing 100 µg/mL hygromycin B (Sigma-Aldrich). The cells were harvested on day 4 post-transfection by pipetting, washed twice with cold 1×PBS, flash frozen with liquid nitrogen, and stored at −80°C for subsequent experiments.

For the immunoprecipitation coupled with western blotting to check the interaction of ORF1p WT or M8 and HERC5 (Figure 3C), HEK293T cells were plated in 10 cm dishes at 3×10^6^ cells/dish density in DMEM. On the next day (day 0), the cells were transfected with 6 µg DNA (4 µg of pJM101/L1.3, pJM101/L1.3FLAG or pALAF008 [L1.3FLAG_M8 (RBM)] and 2 µg of pALAF023 [HERC5]) using 18 µL PEI-MAX-40K in 500 µL Opti-MEM. On the following day (day 1), the medium was replaced with fresh DMEM. On days 2 and 3 post-transfection, the medium was replaced with fresh DMEM containing 100 µg/mL hygromycin B. The cells were harvested on day 4 post-transfection by pipetting, washed twice with cold 1×PBS, flash frozen with liquid nitrogen, and stored at −80°C for subsequent experiments.

For the immunoprecipitation coupled with western blotting to check the RC-L1 RNP formation efficiency (Figure 5F), HEK293T cells were plated in 10 cm dishes at 5×10^6^ cells/dish density in DMEM. On the next day (day 0), the cells were transfected with 10 µg DNA (8 µg of pTMH3 or pTMF3 and 2 µg of pCMV-3Tag-9, pALAF023 [HERC5], pALAF024 [MOV10]) using 30 µL PEI-MAX-40K in 500 µL Opti-MEM. On the following day (day 1), the medium was replaced with fresh DMEM. On days 2 and 3 post-transfection, the medium was replaced with fresh DMEM containing 100 µg/mL hygromycin B. The cells were harvested on day 4 post-transfection by pipetting, washed twice with cold 1×PBS, flash frozen with liquid nitrogen, and stored at −80°C for subsequent experiments.

**Figure 5.**
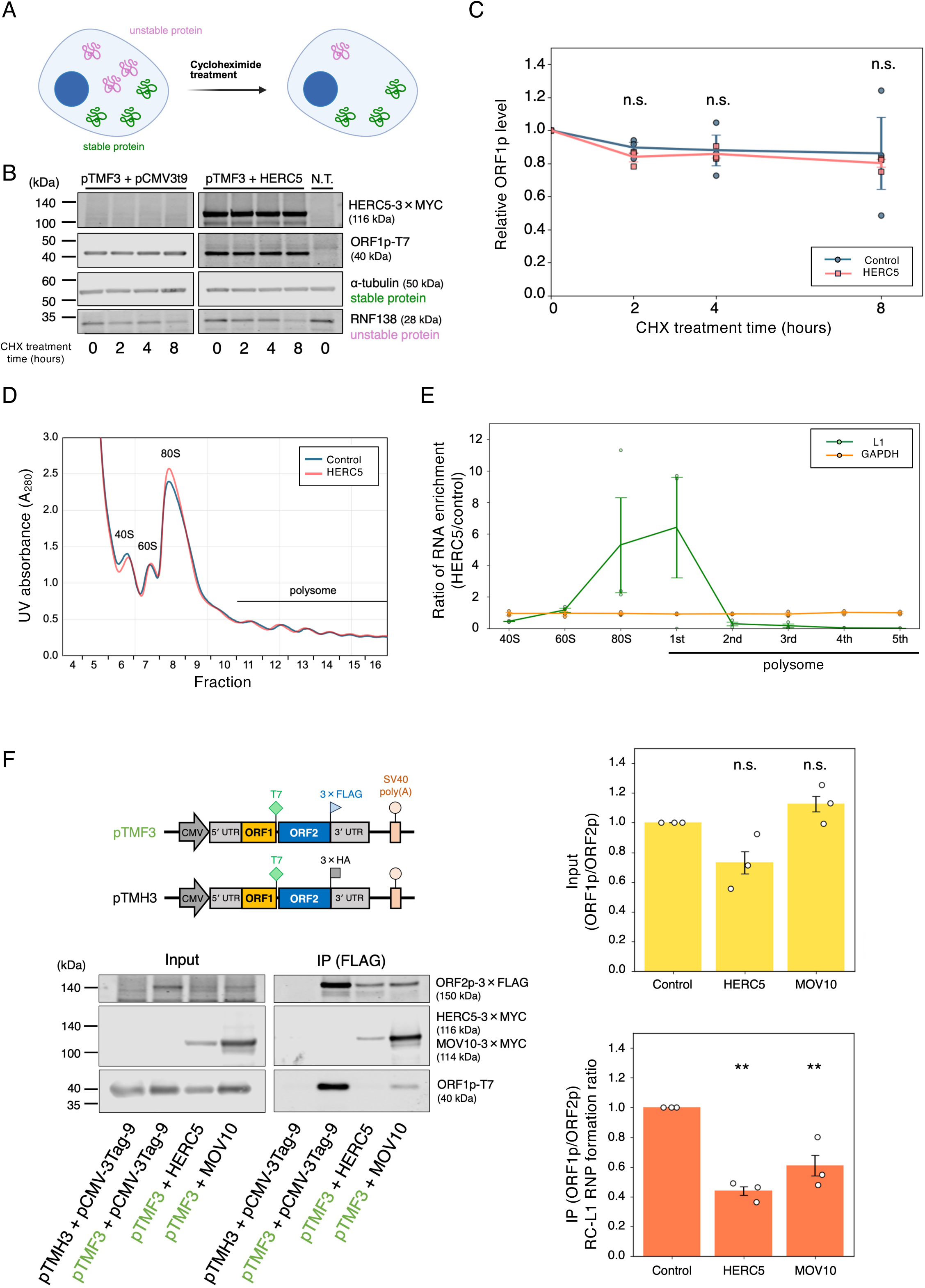
HERC5 reduces L1 translation efficiency and impairs RC-L1 RNP formation. (**A**) Rationale for the cycloheximide (CHX) treatment experiment. CHX treatment stops *de novo* protein synthesis. Unstable proteins (colored magenta) are reduced by CHX treatment due to protein degradation, while stable proteins (colored green) are not. The figure was created with BioRender. (**B**) Protein levels following CHX treatment. HEK293T cells were co-transfected with pTMF3 and either pCMV-3Tag-9 or a HERC5-expressing vector and treated with 50 µg/mL CHX for the indicated times (0, 2, 4, and 8 h). HERC5, ORF1p, α-tubulin, and RNF138 were detected by anti-MYC, anti-T7, anti-α-tubulin, and anti-RNF138 antibodies, respectively. α-tubulin served as a stable protein control, whereas RNF138 served as an unstable protein control. N.T., non-transfected cells. (**C**) Quantification of the remaining ORF1p levels in the control (blue line) and HERC5 (red line) from (B). ORF1p and GAPDH band signal intensities were measured with Empiria Studio. The ORF1p intensities were normalized to GAPDH intensities, and the relative ORF1p levels were calculated. X-axis, CHX treatment time. Y-axis, the relative ORF1p level compared to the ORF1p level at 0 h (both control and HERC5 were set to 1.0, respectively). Each dot represents an independent biological replicate. The *p*-values were calculated using a one-way ANOVA followed by Bonferroni-Holm post-hoc tests; * *p* < 0.05, ** *p* < 0.01, *** *p* < 0.001; n.s.: not significant. (**D**) Polysome profiling analysis of control (blue line) and HERC5 (red line). HEK293T cells were co-transfected with L1 and either pCMV-3Tag-9 (control) or a HERC5 expression vector. Cell lysates were subjected to sucrose gradient centrifugation and separated into the indicated fractions. X-axis, fraction number. Y-axis, UV absorbance at 280 nm, showing the distribution of ribosomal subunits (40S and 60S), monosomes (80S), and polysomes. (**E**) RNA enrichment ratio in polysome fractions. L1 (normalized to GAPDH) and GAPDH RNA levels in each sucrose fraction were measured by RT-qPCR, and the ratio of RNA enrichment (HERC5/control) was calculated. The green and orange lines show L1 and GAPDH RNA ratios, respectively. X-axis, ribosome fraction. Y-axis, ratio of RNA enrichment (HERC5/control). The error bars and *p*-values were calculated as noted in (C). (**F**) RC-L1 RNP formation efficiency with HERC5. HEK293T cells were co-transfected with pTMF3 and either pCMV-3Tag-9, a HERC5-expressing vector, or a MOV10 expression vector. Cells were harvested on 4 days post-transfection, and ORF2p-3×FLAG complexes were immunoprecipitated. Top left: schematic of full-length L1 constructs (pTMF3 and pTMH3). pTMF3 expresses ORF1p tagged with a T7 gene 10 epitope and ORF2p tagged with a 3×FLAG epitope at their carboxyl termini. pTMH3 is similar to pTMF3 but expresses 3×HA epitope-tagged ORF2p and served as a negative control. Bottom left: the input and anti-FLAG IP reactions were analyzed by western blotting. ORF2p and ORF1p were detected by anti-FLAG and anti-T7 antibodies, respectively. HERC5 and MOV10 were detected by an anti-MYC antibody. Right: the input ORF1p/ORF2p ratios (top) and immunoprecipitated ORF1p/ORF2p ratios (bottom), which indicate RC-L1 RNP formation efficiencies. The protein band signal intensities were measured with Empiria Studio. Each ORF1p signal intensity was divided by the respective ORF2p signal intensity, and the resulting ORF1p/ORF2p ratio was calculated. X-axis, name of the transfected constructs. Y-axis, relative ORF1p/ORF2p ratio of input compared to the control (pCMV-3Tag-9, set to 1.0). The error bars and *p*-values were calculated as noted in (C).

The following procedures were performed on ice or at 4°C, unless noted otherwise (Figure 3A, C, and 5F). The antibody-conjugated beads were prepared prior to immunoprecipitation. Ten microliters of Dynabeads Protein G (Invitrogen) were washed twice with 500 µL of 1×PBS containing 0.1% (v/v) Triton X-100 and 0.5% (w/v) Bovine Serum Albumin (BSA) Fraction V, Reagent Grade (MP Biomedicals, California, United States) and incubated with 1 µg of anti-FLAG M2 antibody in 50 µL of 1×PBS containing 0.1% (v/v) Triton X-100 and 0.5% (w/v) BSA for 1.5 h. After incubation, the antibody-conjugated beads were washed twice with 500 µL of 1×PBS containing 0.1% (v/v) Triton X-100 and 0.5% (w/v) BSA. The beads were resuspended in 10 µL/sample of Lysis150 buffer (20 mM Tris-HCl [pH 8.0], 2.5 mM MgCl_2_ [Nacalai Tesque], 150 mM KCl [Nacalai Tesque], and 0.5% [v/v] NP-40/IGEPAL CA-630 [Nacalai Tesque]) containing 1×cOmplete EDTA-free protease inhibitor cocktail, 0.2 mM phenylmethylsulfonyl fluoride (PMSF) (Nacalai Tesque), and 1 mM DTT.

Each cell pellet was lysed in 1 mL Lysis150 buffer containing 1×cOmplete EDTA-free protease inhibitor cocktail, 0.2 mM PMSF, and 1 mM DTT and incubated for 1 h. After incubation, the lysates were centrifuged at 12,000 rpm for 5 min using MDX-310, and the supernatants were collected. Ten microliters of the supernatant were saved as the “input”. The antibody-conjugated beads were added to the samples and incubated for 2–3 h. After incubation, the beads were washed four times with 100 µL of Lysis150 buffer. The ORF1p-FLAG protein complexes were eluted with Lysis150 buffer containing 1×cOmplete EDTA-free protease inhibitor cocktail, 0.2 mM PMSF, 1 mM DTT, and 200 µg/mL 3×FLAG peptide (Sigma-Aldrich) by rotation for 2 h. After elution, 3×SDS sample buffer was directly added to the eluate, which was then boiled at 100°C for 5 min and subjected to western blotting.

For the immunoprecipitation coupled with the RNase treatment (Figure 3B), the beads were washed with 100 µL of Lysis150 buffer after immunoprecipitation. Then, 20 µg/mL of RNase G.S. (Nippon Gene, Tokyo, Japan) was added to the beads in 100 µL of Lysis150 buffer, and the beads were incubated at 37°C for 10 min. The beads were washed four times with 100 µL of Lysis150 buffer. Finally, the beads were resuspended directly in 3×SDS sample buffer, boiled at 100°C for 5 min, and subjected to western blotting.

### Immunoprecipitation for mass spectrometry of the HERC5 complexes

For the HERC5 immunoprecipitation coupled with mass spectrometry (Supplementary Figure S5), HEK293T cells were plated in 10 cm dishes at 2.5×10^6^ cells/dish density in DMEM. Ten dishes were prepared for each sample. On the next day (day 0), the cells were transfected with 10 µg DNA (7 µg of pALAF023 [HERC5-3×MYC] control, pKN002 [HERC5-3×FLAG], pKN005 [HERC5_C994A-3×FLAG]) using 30 µL PEI-MAX-40K in 500 µL Opti-MEM. On the following day (day 1), the medium was replaced with fresh DMEM. On days 2 and 3 post-transfection, the medium was replaced with fresh DMEM containing 100 µg/mL hygromycin B. On day 4 post-transfection, the cells were washed with cold 1×PBS, trypsinized with 0.25% Trypsin-EDTA, and harvested. The cell pellets were flash frozen with liquid nitrogen and stored at −80°C for subsequent experiments.

The following procedures were performed on ice or at 4°C unless noted otherwise. The antibody-conjugated beads were prepared prior to immunoprecipitation. One hundred microliters of Dynabeads Protein G were washed twice with 5 mL of 1×PBS containing 0.1% (v/v) Triton X-100 and 0.5% (w/v) BSA and incubated with 5 µg of anti-FLAG M2 antibody in 500 µL of 1×PBS containing 0.1% (v/v) Triton X-100 and 0.5% (w/v) BSA for 1.5 h. After incubation, the antibody-conjugated beads were washed twice with 500 µL of 1×PBS containing 0.1% (v/v) Triton X-100 and 0.5% (w/v) BSA. The beads were resuspended in 100 µL/sample of Lysis150 buffer containing 1×cOmplete EDTA-free protease inhibitor cocktail, 0.2 mM PMSF, and 1 mM DTT.

Each cell pellet was lysed in 5 mL Lysis150 buffer containing 1×cOmplete EDTA-free protease inhibitor cocktail, 0.2 mM PMSF, and 1 mM DTT and incubated for 1 h. After incubation, the lysates were centrifuged at 12,000 rpm for 5 min using MX-300 (Tomy Seiko, Tokyo, Japan), and the supernatants were collected. Ten microliters of the supernatant were saved as the “input”. The antibody-conjugated beads were added to the samples and incubated for 2–3 h. After incubation, the beads were washed five times with 500 µL of Lysis150 buffer. The HERC5-FLAG complexes were eluted twice in 100 µL of Lysis150 buffer containing 1×cOmplete EDTA-free protease inhibitor cocktail, 0.2 mM PMSF, 1 mM DTT, and 200 µg/mL 3×FLAG peptide by rotating for 2 h each time. After elution, the resultant eluates (200 µL in total) were incubated with 700 µL of acetone (Nacalai Tesque) at −20°C overnight. The protein complexes were precipitated by centrifugation at 15,000 rpm using MX-300, 4°C for 15 min. After the supernatant was removed, the protein pellets were dried at room temperature. Thirty microliters of 1.5×SDS sample buffer was added to the pellets and boiled at 105°C for 5 min. The denatured proteins were stored at −80°C and used for subsequent experiments.

Mass spectrometry analysis was performed at the Proteomics Facility of the Graduate School of Biostudies, Kyoto University. Protein samples were separated by SDS-PAGE and visualized using the PlusOne Silver Staining Kit, Protein (Cytiva, Massachusetts, United States), according to the manufacturer’s protocol. Each gel lane was excised into 18 slices. The silver stain was removed from the gels, and they were subjected to in-gel digestion with sequencing-grade modified trypsin (Promega) to extract peptides. Purified peptides were analyzed by liquid chromatography-tandem mass spectrometry (LC–MS/MS) using a nano-Advance (AMR, Tokyo, Japan) and Q Exactive Plus (Thermo Fisher Scientific) using Xcalibur 3.1 (Thermo Fisher Scientific), Paradigm Home v.2.0.4 R4 B22 (Bruker Daltonics, Massachusetts, United States). For label-free quantification (LFQ) analysis, protein identification was performed using Mascot Server 2.7.0 (Matrix Science; https://www.matrixscience.com/) as a search engine against UniProt Knowledgebase (UniProtKB: https://www.uniprot.org/help/uniprotkb), and protein abundances were calculated using Proteome Discoverer 2.3 (Thermo Fisher Scientific). The Protein Validator Node of Proteome Discoverer 2.3 calculated high (<0.01), medium (0.01≤FDR<0.05), or low (FDR≥0.05) confidence. Both unique and razor peptides were used for protein group assignment. Razor peptides, which are shared among multiple protein groups, were assigned to the group with the highest total peptide count when combined with unique peptides. Triplicate datasets from the 18 gel slices of each sample were grouped, and protein abundance was quantified using the Precursor Ions Quantifier node in Proteome Discoverer 2.3. Abundances were normalized to HERC5 protein levels. The normalized abundances were compared between HERC5 WT and C994A samples to calculate the abundance ratios, whose ratio thresholds were set at a minimum of 0.001 and a maximum of 1000. HERC5 mass spectrometry followed by LFQ was conducted on three independent biological replicates. Statistical analyses were performed using ANOVA followed by Tukey’s Honestly Significant Difference (HSD) post hoc test. The mass spectrometry proteomics data have been deposited to the ProteomeXchange Consortium via the PRIDE partner repository with the identifier PXD068064 and 10.6019/PXD068064 (Token: 4h8ae9WHQpHM).

### GO term analysis

Prior to Gene Ontology (GO) term analysis, proteins were filtered based on the following criteria to identify proteins commonly associated with both HERC5 WT and C994A mutant:

#### 1. LFQ abundance ratios

In the LFQ analysis, 1,824 proteins with abundance ratios between 0.5 and 2.0 and high FDR confidence scores were retained.

#### 2. Mascot prot_matches

Proteins were filtered by comparing Mascot prot_matches across all biological replicates. Only proteins that met both of the following criteria were retained.

- Proteins showing a difference of more than 4 in Mascot prot_matches between HERC5 WT-FLAG and HERC5-MYC (negative control).
- Proteins showing a difference of more than 4 in Mascot prot_matches between HERC5 C994A-FLAG and HERC5-MYC (negative control).

A total of 428 proteins were selected from the Mascot prot_matches selection. Of those, 316 proteins, which satisfied both the LFQ abundance ratios and Mascot prot_matches criteria, were subjected to GO term analysis using the Database for Annotation, Visualization, and Integrated Discovery (DAVID) analysis (https://davidbioinformatics.nih.gov/) to obtain the “Biological Processes” GO terms (85, 86).

### Primary antibodies used in this study

The following list provides information on the primary antibodies used in this study, including the host species, clonality, target antigen, dilution factor, supplier, catalog number, and Research Resource Identifier (RRID).

Mouse polyclonal anti-HERC5 antibody (1/500), (Abnova, H00051191-A01, RRID: AB_894114)

Mouse monoclonal anti-GAPDH antibody (1/1,000), (Millipore, MAB374, RRID: AB_2107445)

Mouse monoclonal anti-FLAG M2 antibody (1/1,000), (Sigma-Aldrich, F1804, RRID: AB_262044)

Rabbit polyclonal anti-FLAG antibody (1/1,000), (Millipore, F7425-2MG, RRID: AB_439687)

Mouse monoclonal anti-MYC antibody (1/1,000), (Cell Signaling Technology, 9B11, RRID: AB_331783)

Rabbit polyclonal anti-T7-tag antibody (1/1,000), (Cell Signaling Technology, D9E1×, RRID: AB_2798161)

Mouse monoclonal anti-ORF1p (4H1) antibody (1/1,000), (Millipore, MABC1152, RRID: AB_2941775)

Mouse monoclonal anti-α-tubulin antibody (1/1,000), (Sigma-Aldrich, T9026, RRID: AB_477593)

Rabbit polyclonal anti-RNF138 antibody (1/1,000), (Sigma-Aldrich, SAB4502131, RRID: AB_10762596)

Goat polyclonal anti-luciferase antibody (1/1,000), (Promega, G7451, RRID: AB_430862)

### Secondary antibodies used in this study

The following list provides information on the secondary antibodies used in this study, including the host species, clonality, target antigen, dilution factor, supplier, catalog number, and Research Resource Identifier (RRID).

Donkey polyclonal anti-rabbit IRDye 680RD-conjugated antibody (1/10,000), (LI-COR, 925-68073, RRID: AB_2716687)

Donkey polyclonal anti-mouse IRDye 680RD-conjugated antibody (1/10,000), (LI-COR, 925-68072, RRID: AB_2814912)

Donkey anti-rabbit IRDye 800CW-conjugated antibody (1/10,000), (LI-COR, 926-32213, RRID: AB_621848)

Donkey anti-mouse IRDye 800CW-conjugated antibody (1/10,000), (LI-COR, AB_621847) 926-32212, RRID: AB_621847)

Donkey anti-goat IRDye 800CW-conjugated antibody (1/10,000), (LI-COR, AB_2687553) 926-32214, RRID: AB_2687553)

### Immunofluorescence

For the observation of HERC5 and ORF1p localization (Supplementary Figure S2B), HEK293T cells were plated in 6-well plates at 1×10^5^ cells/well density in DMEM. On the next day (day 0), the cells were transfected with 1 µg DNA (0.5 µg of pJM101/L1.3 or pJM101/L1.3FLAG and 0.5 µg of pCMV-3Tag-9, pALAF023 [HERC5 WT], pALAF062 [HERC5 ΔRLD], pALAF061 [HERC5 ΔHECT], pALAF060 [HERC5 C994A]) using 3 µL FuGENE HD in 100 µL Opti-MEM according to the manufacturer’s protocol. On the following day (day 1), the medium was replaced with fresh DMEM. On day 2, the cells were trypsinized with 0.25% Trypsin-EDTA and 1×10^5^ cells were replated to 18 mm glass coverslips (Matsunami Glass, Osaka, Japan) coated with 50 µg/mL poly-L-lysine (Sigma-Aldrich) in 12-well plate (BioLite, New York, United States). Approximately 24 h later, cells were washed with cold 1×PBS and fixed with 4% paraformaldehyde (Electron Microscopy Sciences, Pennsylvania, United States) at room temperature for 10 min. After fixation, cells were washed with cold 1×PBS containing 10 mM glycine for 5 min.

The cells were permeabilized with 0.5% (v/v) Triton X-100 for 3 min. The cells were washed twice with cold 1×PBS containing 10 mM glycine for 5 min. The primary antibodies (1/500 dilution in PBST [0.1% (v/v) Tween-20 in 1×PBS]) containing 0.5% (v/v) Normal Donkey Serum (NDS, Sigma-Aldrich) and 10 mM glycine were incubated on the coverslip for 45 min at room temperature. The cells were washed three times with 1×PBS containing 10 mM glycine for 5 min. The secondary antibodies (1/1,000 dilution in PBST) containing 0.5% (v/v) NDS, 10 mM glycine, and 1 µg/mL 4′, 6-diamidino-2-phenylindole (DAPI) (Sigma-Aldrich) were incubated on the coverslip for 45 min at room temperature. The cells were washed three times with cold 1×PBS containing 10 mM glycine for 5 min, followed by a final rinse with Milli-Q. The glass coverslips were mounted on slides with 3 µL VECTASHIELD (Vector Laboratories, Burlingame, CA, United States).

Images were captured by a fluorescence microscope BZ-X800 (Keyence, Osaka, Japan) and visualized with the BZ-X800 Analyzer version 1.1.2.4 (Keyence). FLAG-tagged ORF1p was probed with an Alexa 488-conjugated antibody (Thermo Fisher Scientific) and visualized with the GFP channel. MYC-tagged HERC5 proteins were probed with an Alexa 568-conjugated antibody (Thermo Fisher Scientific) and visualized with the TRITC channel. The DAPI signal was visualized with the DAPI channel.

### Cycloheximide chase assay

For the cycloheximide chase assay (Figure 5B and C), HEK293T cells were plated in 6 cm dishes at 4×10^5^ cells/dish density in DMEM. On the next day (day 0), the cells were transfected with 2 µg DNA (1 µg of pTMF3 and 1 µg of pCMV-3Tag-9 or pALAF023 [HERC5]) using 6 µL FuGENE HD in 200 µL Opti-MEM according to the manufacturer’s protocol. On the following day (day 1), the medium was replaced with fresh DMEM without antibiotics. On day 2 post-transfection, cells were harvested at 0, 2, 4, and 8 h after treatment with 50 µg/mL cycloheximide (CHX) (Nacalai Tesque). HEK293T cells were then harvested by pipetting and washed twice with cold 1×PBS. The cell pellets were flash frozen with liquid nitrogen and stored at −80°C. The protein levels were analyzed by western blotting as described in the “Western blots” section.

### Polysome profiling

For the polysome profiling to check L1 translation efficiency (Figure 5D and E), HEK293T cells were plated in two 10 cm dishes at 3×10^6^ cells/dish density in DMEM. On the next day (day 0), the cells were transfected with 6 µg DNA (4 µg of pTMF3 and 2 µg of pCMV-3Tag-9 or pALAF023 [HERC5]) using 18 µL PEI-MAX-40K in 500 µL Opti-MEM. On days 1 and 2, the medium was replaced with fresh DMEM. On day 3 post-transfection, cells were incubated for 30 min in DMEM containing 100 µg/mL CHX at 37°C, harvested by pipetting, and washed twice with cold 1×PBS containing 100 µg/mL CHX. The collected cells were lysed with lysis buffer (20 mM HEPES-NaOH [pH 7.5] [Nacalai Tesque], 2.5 mM MgCl_2_, 150 mM NaCl, and 1% [v/v] Triton X-100) containing 1×cOmplete EDTA-free protease inhibitor cocktail, 1×PhosSTOP phosphatase inhibitor cocktail (Roche Diagnostics), 1 mM DTT, and 100 µg/mL CHX. The cells were incubated in the buffer for 5 min and centrifuged at 12,000 rpm at 4°C for 5 min using MDX-310. The resultant cell lysates were loaded onto 15–60% (w/v) sucrose (Nacalai Tesque) gradient solution containing 20 mM HEPES-NaOH (pH 7.5), 2.5 mM MgCl_2_, 150 mM NaCl, and 1 mM DTT, and then centrifuged at 30,000 rpm at 4°C for 2.5 h using SW41Ti rotor (Beckman Coulter, California, United States). After centrifugation, each fraction (500 µL) was collected from the top to the bottom of the gradient using Triax Flow Cell (FC-2) (Biocomp Instruments, British Columbia, Canada), monitoring UV absorbance at 280 nm. Two hundred fifty microliters of each sucrose fraction was mixed with 750 µL TRIzol (Invitrogen) and 180 µL chloroform (Nacalai Tesque), and shaken vigorously for 15 sec. The tubes were incubated at room temperature for 5 min and centrifuged at 12,000 rpm at 4°C for 15 min using MDX-310. Then, 360 µL of the upper aqueous layer was collected and mixed with 400 µL isopropanol (Nacalai Tesque) and 400 µL of 0.8 M citric acid (Nacalai Tesque) solution containing 1.2 M NaCl to remove sucrose. The solution was shaken well and incubated at room temperature for 5 min, and RNA was pelleted by centrifugation at 12,000 rpm at 4°C for 30 min using MDX-310. After centrifugation, the pellets were washed with 180 µL of 75% (v/v) ethanol and centrifuged again at 12,000 rpm at 4°C for 5 min using MDX-310. Ethanol was removed, and the pellets were dried at room temperature, then resuspended in 30 µL RNase-free water. RT-qPCR was performed as described in the RT-qPCR section.

### Statistical analysis

One-way ANOVA followed by Bonferroni-Holm post hoc tests was performed using the online web statistical calculators ASTATSA 2016 (https://astatsa.com/). The number of biological replicates is indicated in the figure legends. The error bars represent the mean ± the standard error of the mean (SEM) of independent biological replicates. The *p*-values of each pair were indicated in the figure legends. * indicates *p* < 0.05, ** indicates *p* < 0.01, *** indicates *p* < 0.001, and n.s. indicates not significant.

## Results

### HERC5 inhibits L1 retrotransposition independently of ISGylation

We previously showed that HERC5 interacts with ORF1p and inhibits L1 retrotransposition in HeLa-JVM and HEK293T cells (52). To examine the contribution of each domain to HERC5-mediated inhibition of L1 retrotransposition, we generated HERC5 expression constructs in the pCMV-3Tag-9 vector, including HERC5 wild-type (WT), an RLD domain-deleted mutant (ΔRLD), a HECT domain-deleted mutant (ΔHECT), and an ISGylation-defective catalytic mutant (C994A) (Figure 1A). We investigated the effects of these mutants on L1 retrotransposition by co-transfecting HEK293T cells with an empty vector (control), HERC5 WT, ΔRLD, ΔHECT, C994A, or MOV10 (positive control)-expressing plasmid together with cepB-gfp-L1.3, which expresses an engineered human L1.3 containing the *mEGFPI* retrotransposition indicator cassette (Figure 1B, see Materials and Methods) (52, 74). L1 retrotransposition efficiencies were determined by measuring the percentage of EGFP-positive cells using flow cytometry (Supplementary Figure S1A). Overexpression of HERC5 WT and C994A similarly reduced L1 retrotransposition efficiency by ∼80% compared to the control, suggesting that HERC5 inhibits L1 retrotransposition independently of its ISGylation activity (Figure 1C). ΔHECT also reduced L1 retrotransposition efficiency by ∼20%, albeit much less than WT, whereas ΔRLD had no significant effect, suggesting that the RLD domain is indispensable for L1 inhibition in HEK293T cells. We also examined expression levels of HERC5 and its mutants. When HERC5 was expressed from pCMV-3Tag-9, C994A protein levels were comparable to WT, suggesting that the point mutation did not affect protein stability; however, ΔRLD and ΔHECT showed lower protein levels than WT (Figure 1D). To validate our results, we expressed HERC5 from a different vector backbone (pEBNA) and assessed the effects on L1 retrotransposition. In this context, HERC5 WT and C994A reduced L1 retrotransposition efficiency by ∼90%, and ΔHECT by ∼85%, whereas ΔRLD did not significantly reduce L1 retrotransposition efficiency (Supplementary Figure S1B, top). Of note, the pEBNA vector expressed ΔRLD and ΔHECT proteins at levels comparable to WT (Supplementary Figure S1B, bottom), reinforcing that the RLD domain is indispensable for L1 inhibition and that the HECT domain is not required for inhibition when ΔHECT is expressed at sufficient levels.

We observed a similar trend of L1 inhibition by HERC5 in HeLa-JVM cells when HERC5 was overexpressed from pCMV-3Tag-9 (Supplementary Figure S1C): ΔRLD and ΔHECT did not significantly affect L1 retrotransposition, while HERC5 WT and C994A similarly reduced L1 retrotransposition efficiency by ∼50% compared to the control, suggesting that ISGylation-independent inhibition by HERC5 is conserved across cell types.

To test whether endogenous HERC5 inhibits L1 retrotransposition, we knocked down HERC5 using small interfering RNA (siRNA) or short hairpin RNA (shRNA) in HEK293T and HeLa-JVM cells, respectively. HERC5 siRNA-treated HEK293T cells showed a slight, though not significant, increase in retrotransposition efficiency compared to the control knockdown cells (Figure 1E, left). We speculate that this may be due to low endogenous HERC5 levels, and therefore, the knockdown had minimal effect on retrotransposition. To increase endogenous HERC5 levels, we treated HEK293T cells with interferon-α (IFN-α). As expected (48), IFN-α treatment increased HERC5 levels and reduced L1 retrotransposition efficiency (Figure 1E and F; IFN-α [−] vs [+]). Comparing the retrotransposition efficiencies of the IFN-treated cells to those of the untreated cells (IFN-treated/untreated ratio) revealed that HERC5 knockdown modestly but significantly increased L1 retrotransposition efficiency in IFN-α-treated cells (Figure 1E, right; siHERC5 vs siControl; see also Figure 1F). In HeLa-JVM cells, HERC5 knockdown increased L1 retrotransposition ∼3-fold compared to the control knockdown (Supplementary Figure S1D). A side-by-side comparison suggested that the endogenous HERC5 levels are higher in HeLa-JVM than in HEK293T cells (Supplementary Figure S1E), which may explain the minimal effect of HERC5 knockdown in untreated HEK293T cells. Alternatively, HERC5 may be the dominant ISG that potentially inhibits L1 in HeLa-JVM cells, as it was significantly enriched in our previous mass-spectrometry analysis of L1 RNPs (52).

### HERC5 reduces ORF1p but not L1 RNA levels

To elucidate the mechanism by which HERC5 inhibits L1 retrotransposition, we examined its effects on L1 RNA and ORF1p. HEK293T cells were co-transfected with a full-length L1 expression plasmid (pTMF3) and either an empty vector or a HERC5 expression plasmid (Figure 2A). RT-qPCR analysis, using a primer pair that specifically recognizes the *T7 gene 10* epitope-tag sequence in the L1 plasmid, revealed no significant change in L1 RNA levels upon HERC5 overexpression (Figure 2B). In contrast, as previously reported (52), HELZ2 overexpression led to a ∼80% reduction in L1 RNA levels compared to the control. This suggests that HERC5 does not significantly affect L1 RNA levels.

Next, we analyzed ORF1p levels with HERC5 overexpression by western blotting and flow cytometry in HEK293T cells. The western blot analysis demonstrated that HERC5 overexpression markedly decreased ORF1p levels compared to the control (Figure 2C). Using an EGFP-tagged ORF1p construct (pVan583) and flow cytometry, we found that HERC5 reduced both the percentage of EGFP-positive cells and the median fluorescence intensity (Figure 2D). In contrast, these effects were not observed when EGFP alone was expressed (pKN033) (Figure 2E), suggesting that HERC5 specifically downregulates ORF1p. Consistently, HERC5 knockdown increased ORF1p levels with or without IFN-α treatment (Supplementary Figure S2A). Taken together, these data strongly support the conclusion that HERC5 inhibits L1 retrotransposition primarily by lowering ORF1p levels rather than by decreasing L1 RNA levels.

### The RLD domain is required for ORF1p reduction

To identify the HERC5 domain responsible for reducing ORF1p, we measured ORF1p-EGFP levels by flow cytometry in HEK293T cells co-transfected with the L1 plasmid (pVan583) and either HERC5 WT or the mutant constructs. HERC5 WT significantly reduced both the percentage of EGFP-positive cells and the median fluorescence intensity, consistent with the previous results (Figure 2F). Conversely, ΔRLD did not show a significant reduction in the percentage of EGFP-positive cells relative to the control (Figure 2F, left). Although the median EGFP intensity in ΔRLD-transfected cells was lower than that in the control, the reduction was smaller than that with HERC5 WT or the other mutants (Figure 2F, right), suggesting that the RLD domain is important for ORF1p reduction. Notably, both ΔHECT and C994A mutants retained the ability to reduce both the percentage and median intensity to levels comparable to WT, suggesting that the HECT domain and its ISGylation E3 ligase activity are dispensable for ORF1p reduction.

### Deletion of the RLD domain alters HERC5 subcellular localization

We analyzed the subcellular localization of HERC5 and its mutants, and their effects on ORF1p expression, using immunofluorescence staining. HEK293T cells were co-transfected with a full-length L1 plasmid expressing FLAG-tagged ORF1p (pJM101/L1.3FLAG) and either the empty vector, HERC5 WT, or its mutants. In the control, cytoplasmic ORF1p foci were observed, consistent with previous reports (23, 47, 52). HERC5 WT expression markedly reduced the ORF1p signal, and it was localized exclusively in the cytoplasm. In contrast, ΔRLD was detected in both the nucleus and cytoplasm, and the ORF1p signals were more evident in cells expressing ΔRLD, further suggesting that the RLD domain is important for reducing ORF1p levels (Supplementary Figure S2B). ΔHECT and C994A mutants, similar to WT, also localized in the cytoplasm and reduced ORF1p expression. These results imply that the RLD domain contributes to HERC5 cytoplasmic localization and is important for its access to L1 RNPs. In a previous study (52), ORF1p foci were detectable in ∼50% of transfected cells even under HERC5 overexpression; in contrast, here we did not clearly detect ORF1p foci by immunofluorescence. A likely explanation for this discrepancy is a cell line difference: the previous study used HeLa-JVM cells, whereas we used HEK293T cells in this study. Because HERC5-mediated L1 inhibition seems to be more robust in HEK293T compared to HeLa-JVM (Figure 1C vs Supplementary Figure S1C), ORF1p levels in HEK293T were likely reduced below the detection threshold, precluding visualization of ORF1p foci.

### The RLD domain interacts with L1 RNA independently of ORF1p

We next performed co-immunoprecipitation of FLAG-tagged ORF1p with HERC5 variants to determine which domain of HERC5 is important for the interaction with ORF1p (Figure 3A, top). HERC5 WT, ΔHECT, and C994A were co-immunoprecipitated with FLAG-tagged ORF1p, whereas ΔRLD was not, indicating that the RLD domain is essential for the ORF1p-HERC5 interaction (Figure 3A and Supplementary Figure S3A).

Because ORF1p together with ORF2p binds L1 RNA in cis to form L1 RNPs (20–23), we asked whether HERC5 interacts with ORF1p directly or through L1 RNA. To address this, we performed co-immunoprecipitation with FLAG-tagged ORF1p followed by RNase A treatment, which degrades single-stranded RNA. RNase A treatment substantially reduced the ORF1p-HERC5 interaction, suggesting that RNA mediates the interaction (Figure 3B). To further confirm the association of HERC5 with L1 RNA, we used the RNA-binding-deficient ORF1p mutant (hereafter referred to as RBM [RNA-binding mutant]), which contains three point mutations (R206A, R210A, and R211A) (Figure 3C, top) (15, 52). Because RBM is unable to bind L1 RNA, testing its interaction with HERC5 could provide insight into whether HERC5 primarily interacts with ORF1p or L1 RNA. Despite the weaker pull-down efficiency, RBM ORF1p binding to HERC5 was notably reduced compared to the WT ORF1p-HERC5 interaction (Figure 3C and Supplementary Figure S3B). The RBM ORF1p-FLAG immunoprecipitation efficiency was lower than that of the WT ORF1p-FLAG, likely because RBM ORF1p fails to bind L1 RNA and thus does not recruit other ORF1p molecules via RNA. These immunoprecipitation results suggest that HERC5 primarily interacts with L1 RNA rather than directly with ORF1p.

To test whether HERC5 associates with L1 RNA in a sequence-dependent manner, we used synthetic human L1 (L1 ORFeus)-expressing plasmids derived from pDA093_tet-ORFeus_SBtet-RN (43). The L1 ORFeus contains codon-optimized RNA that differs by ∼25% from the human L1.3 *ORF* sequences at the nucleotide level but encodes ORF1p and ORF2p, which are ∼99% identical to those of L1.3 at the amino acid level. We measured both L1 ORFeus retrotransposition efficiency and its protein levels. HERC5 reduced native L1.3 retrotransposition by ∼80% compared to the control but reduced L1 ORFeus retrotransposition by ∼40%, suggesting that L1 inhibition by HERC5 is partly dependent on the RNA sequence (Figure 3D). Consistently, HERC5 strongly reduced L1.3-derived ORF1p, whereas L1 ORFeus-derived ORF1p was largely unaffected (Figure 3E). These results suggest that HERC5 suppresses L1 retrotransposition in part by targeting an L1.3-specific RNA sequence (see “Discussion”). However, because HERC5 still reduced L1 ORFeus retrotransposition efficiency by ∼40%, HERC5 may also employ sequence-independent mechanisms to inhibit L1 retrotransposition.

We also tested whether HERC5 targets an L1 RNP-specific structure by comparing levels of WT and RBM ORF1p, the latter of which is thought to be unable to form L1 RNPs due to a loss of RNA binding (15). Intriguingly, as reported previously (52), HERC5 reduced RBM ORF1p to a level comparable to WT ORF1p (Supplementary Figure S3C). This result indicates that L1 RNP formation is not essential for HERC5 to recognize L1. Together, these data support the hypothesis that HERC5 interacts with L1 RNA independently of ORF1p, and that the RLD domain is necessary for this interaction.

### HERC5 regulates ORF1p and its downstream ORF2p

We further investigated which part of L1 is recognized by HERC5 using plasmids expressing different regions of L1. First, to test whether the L1 *ORF* sequences are targeted by HERC5, we replaced the L1 coding region with the luciferase *ORF*. However, we found that the luciferase protein level was not reduced by HERC5, suggesting that the L1 *ORF*s are important for HERC5 activity (Supplementary Figure S4A). In addition, HERC5 reduced ORF1p levels expressed from an L1 construct without 5′ UTR (pTMF3_Δ5UTR) to a level comparable to ORF1p from the full-length L1 construct (pTMF3), suggesting that 5′ UTR is not targeted. Next, we co-expressed HERC5 with monocistronic ORF1p, monocistronic ORF2p, or full-length bicistronic L1. HERC5 reduced monocistronic ORF1p but not monocistronic ORF2p levels (Figure 4A and B). In contrast, ORF2p from the bicistronic L1 construct was reduced together with ORF1p (Figure 4C). These results suggest that HERC5 specifically recognizes the L1.3 *ORF1* sequence for inhibition and consequently affects the downstream ORF2p expressed from the same RNA.

To further test this hypothesis, we investigated the effects of HERC5 on *Alu* retrotransposition in the presence or absence of ORF1p. *Alu* is a non-autonomous retrotransposon comprising ∼11% of the human genome (3). Unlike L1, *Alu* retrotransposition does not strictly require L1-derived ORF1p, but depends on ORF2p (81, 87, 27, 88). pKN040 expresses *Alu* and the full-length bicistronic L1, while pTM489 expresses *Alu* and monocistronic ORF2p lacking ORF1p (Figure 4D and E). We then measured *Alu* retrotransposition efficiencies with these two plasmids in the HeLa-HA cell line, which is permissive for *Alu* retrotransposition (71, 72). HERC5 reduced *Alu* retrotransposition driven by the bicistronic L1 (pKN040) by ∼50% relative to the control (Figure 4D). In contrast, HERC5 had no detectable effect on *Alu* retrotransposition induced by monocistronic ORF2p (pTM489) (Figure 4E). These findings strongly support our hypothesis that HERC5 specifically targets the *ORF1* sequence and reduces its protein level, and that ORF2p is affected only when expressed downstream of *ORF1*. We also confirmed that HERC5-mediated L1 inhibition was conserved in HeLa-HA cells (Supplementary Figure S4B).

### HERC5 reduces L1 translation efficiency

Our results suggest that HERC5 interacts with L1 RNA and modulates ORF1p levels. We hypothesized that HERC5 regulates ORF1p levels either by promoting protein degradation or by suppressing translation. To distinguish these two possibilities, we first treated HEK293T cells with cycloheximide (CHX), a global mRNA translation inhibitor, to block *de novo* protein synthesis and monitored the remaining ORF1p levels with or without HERC5 overexpression (Figure 5A). After CHX treatment, no significant difference in ORF1p levels was observed between conditions with and without HERC5 overexpression, suggesting that HERC5 does not measurably affect ORF1p stability and may instead inhibit its translation (Figure 5B and C). We therefore performed polysome profiling to examine the translation efficiency of L1. HEK293T cells were co-transfected with HERC5 and L1 constructs and treated with CHX to arrest ribosomal elongation on mRNA prior to cell lysis. Cell lysates were then centrifuged in 15–60% (w/v) sucrose density gradients to separate monosome and polysome fractions (Figure 5D). For each fraction, we calculated the enrichment ratios (HERC5/control) for L1 and GAPDH RNAs. Under the HERC5 overexpression condition, L1 RNA accumulated in the 80S and light polysome (first polysome) fractions and was reduced in the heavier polysome fractions (Figure 5E). In contrast, the enrichment ratios for GAPDH RNA were unchanged by HERC5, suggesting that HERC5 selectively reduces L1 translation efficiency, thereby reducing ORF1p levels.

We also performed immunoprecipitation of FLAG-tagged HERC5 WT and C994A followed by LC–MS/MS analysis to identify potential cofactors (Supplementary Figure S5A). FLAG-tagged HERC5 WT and C994A were efficiently immunoprecipitated with anti-FLAG antibody (Supplementary Figure S5B). The analysis revealed that both HERC5 WT and C994A co-purified with multiple translation-related proteins (Supplementary Figure S5C and Table S1), consistent with a previous report showing that HERC5 interacts with polysomes via its RLD domain (64). Collectively, these results suggest that HERC5 potentially forms a complex with translation-associated proteins that dampen L1 translation. Nonetheless, further studies are required to elucidate the cofactors involved and to determine how HERC5 specifically targets L1 RNA for translational regulation.

### HERC5 reduces the efficiency of RC-L1 RNP formation

We next asked whether HERC5 inhibits the formation of retrotransposition-competent L1 RNPs (RC-L1 RNPs) by depleting ORF1p. To obtain RC-L1 RNPs that contain both ORF1p and ORF2p, we purified the FLAG-tagged ORF2p using an anti-FLAG antibody and analyzed the ORF1p and ORF2p levels. In the input lysates, the ORF1p/ORF2p ratio was not significantly altered by HERC5 overexpression, suggesting that HERC5 reduces both ORF1p and ORF2p levels to a similar extent. In contrast, in the immunoprecipitated RC-L1 RNP fractions, HERC5 reduced the ORF1p/ORF2p ratio by ∼60%, suggesting that HERC5 also decreases ORF1p incorporation into RC-L1 RNPs in addition to translational repression (see “Discussion”).

### HERC5 inhibits human and mouse retrotransposons but not a zebrafish retrotransposon

To examine whether the retrotransposition inhibition by HERC5 extends to other retrotransposons, we tested two mouse L1 elements from different subfamilies (TG_F_21, G_F_ subfamily, and ORFeus-Mm, a synthetic mouse T_F_ subfamily) and a zebrafish L2 element (ZfL2-2). In mouse TG_F_21 and ORFeus-Mm retrotransposition assays, HERC5 significantly reduced retrotransposition efficiencies compared to controls (Figure 6A and B). In contrast, HERC5 did not reduce the zebrafish ZfL2-2 retrotransposition efficiency (Figure 6C). This may reflect the lack of *ORF1* in ZfL2-2, which contains a single *ORF* encoding EN and RT domains similar to L1 *ORF2* (78, 89) (Figure 6D). These data further support the notion that HERC5 specifically targets the L1 *ORF1* sequence and that its inhibitory effect is species- and element-specific.

**Figure 6.**
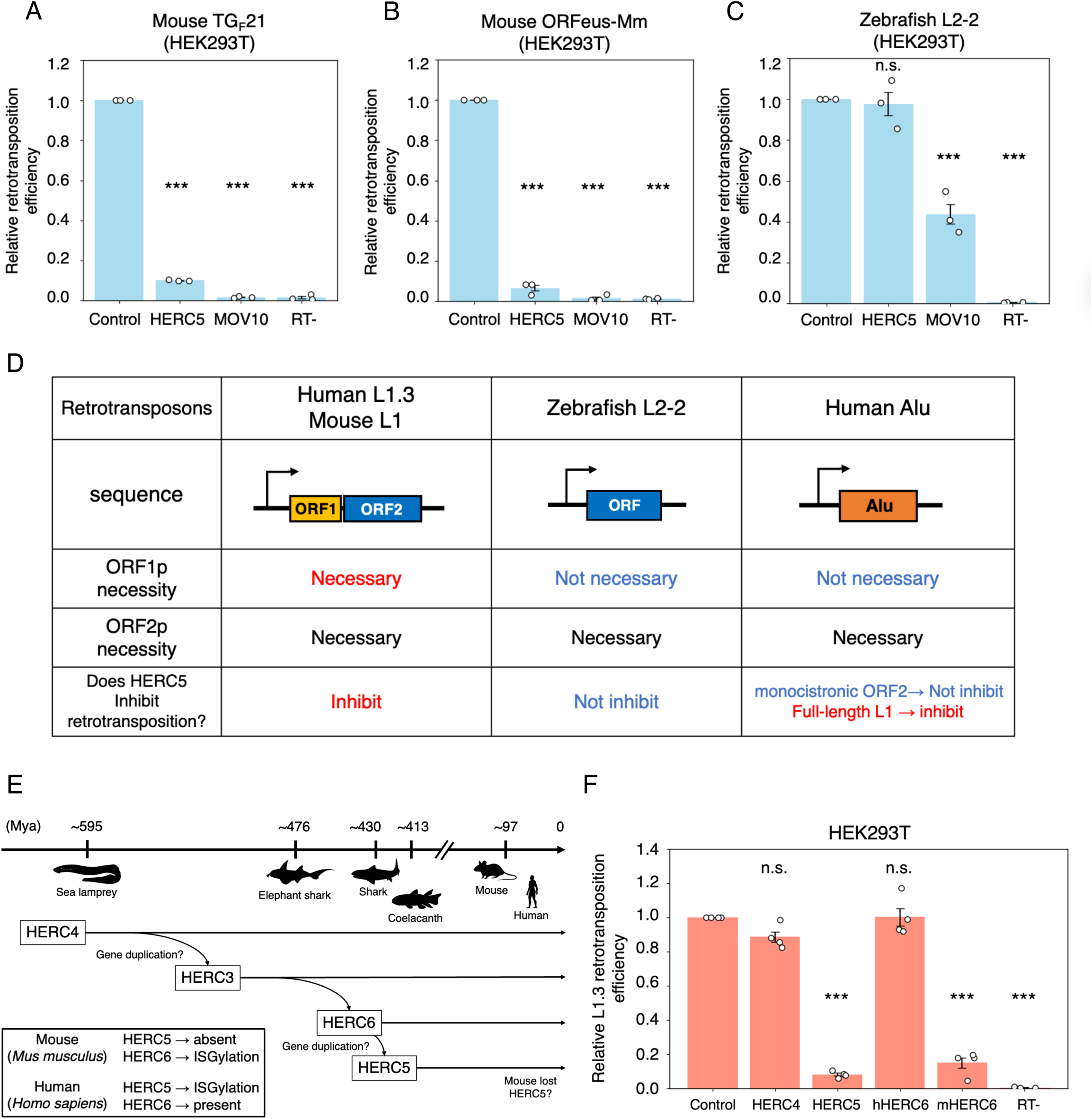
The small HERC family may have acquired the L1-inhibitory mechanism during evolution. (**A–C**) Differential effects of HERC5 on non-human LINE retrotransposition. To measure retrotransposition efficiency, HEK293T cells were co-transfected with either mouse TG_F_21 (A), mouse ORFeus-Mm (B), or zebrafish L2-2 (C) together with either the control vector, a HERC5-expressing vector, or a MOV10-expressing vector. Cells were selected with puromycin (1 µg/mL), and the percentage of EGFP-positive cells was determined by flow cytometry. Cells co-transfected with cep99-gfp-L1.3RT(-) intronless served as transfection-normalization controls. X-axis, name of the transfected constructs. Y-axis, relative retrotransposition efficiency compared to the control (pCMV-3Tag-8-Barr, set to 1.0). Each dot represents an independent biological replicate. The *p*-values were calculated using a one-way ANOVA followed by Bonferroni-Holm post-hoc tests; *** *p* < 0.001; n.s.: not significant. (**D**) Summary of HERC5 effects on retrotransposition. A schematic of retrotransposon structures is shown. Requirements for ORF1 and ORF2p, and whether HERC5 inhibits each element, are indicated for different retrotransposons (human L1.3, mouse L1, zebrafish L2-2, and human *Alu*). (**E**) Schematic timeline of the estimated divergence times of small HERC paralogs. HERC4 is predicted to have emerged earliest (>595 million years ago, Mya), HERC3 to have arisen >∼476 Mya, HERC6 ∼430 Mya, and the HERC5 paralog ∼413 Mya. Divergence times are based on the phylogenetic analysis in (91). The silhouette images of organisms were downloaded from PHYLOPIC version 2.0 (https://www.phylopic.org/). Mice lack HERC5 but have HERC6, which has the E3 ligase activity of ISGylation. Humans have both HERC5 and HERC6; only HERC5 has the activity. (**F**) L1 retrotransposition assay with the small HERC family. HEK293T cells were co-transfected with an L1-expressing construct (cepB-gfp-L1.3) and a member of the small HERC family (HERC4, HERC5, hHERC6, and mHERC6). Cells were selected with blasticidin (10 µg/mL), and the percentage of EGFP-positive cells was measured by flow cytometry. The RT mutant served as a control. X-axis, name of the transfected constructs. Y-axis, relative L1 retrotransposition efficiency compared to the control (pCMV-3Tag-9, set to 1.0). The error bars and *p*-values were calculated as noted in (A).

### Human HERC5 may have acquired an inhibitory mechanism against L1 during evolution

The human small HERC family consists of four members: HERC3, HERC4, HERC5, and HERC6 (90, 91). We examined whether other small HERC family proteins can suppress human L1 retrotransposition. Phylogenetic analysis of the small HERC family showed that HERC4 is the most ancient paralog, followed by HERC3, HERC6, and HERC5 (91). The small HERC family is suggested to have evolved more than 595 million years ago (Mya); HERC3 may have arisen by duplication of HERC4 between ∼476 and 595 Mya, and HERC6 and HERC5 may have subsequently emerged ∼430 Mya and ∼413 Mya, respectively (Figure 6E) (91). The retrotransposition assay showed that HERC4 and HERC6 did not inhibit L1 retrotransposition, whereas HERC5 significantly inhibited it (Figure 6F). This suggests that L1 suppression activity is unique to HERC5 among human small HERC family proteins.

We also tested whether mouse HERC6 (mHERC6) can inhibit human L1 retrotransposition because, unlike in humans, mHERC6 functions as the ISGylation E3 ligase instead of HERC5 (92). It has been suggested that mice have lost HERC5, but the ISGylation E3 ligase activity is conserved in HERC6 (Figure 6E) (92). Human HERC6 (hHERC6) did not inhibit human L1 retrotransposition; however, mHERC6 efficiently inhibited human L1 retrotransposition (Figure 6F). This result suggests that mHERC6 acquired L1 suppression activity after the human–mouse divergence. We propose that the small HERC family may have independently evolved mechanisms to restrict L1 retrotransposition (see “Discussion”).

## Discussion

HERC5 has been reported to exert antiviral activity against many viruses, primarily through an ISGylation-dependent mechanism (55). To date, several reports have suggested ISGylation-independent activities of HERC5, including inhibition of HIV-1 and Ebola virus; however, the detailed underlying mechanisms remain poorly understood (69, 70). In this study, we demonstrate that HERC5 inhibits retrotransposition of endogenous retroelements independently of ISGylation through previously unreported mechanisms involving suppression of translation and inhibition of RC-L1 RNP formation by targeting the L1 ORF1 RNA sequence.

We show that the RLD domain of HERC5 is required for this ISGylation-independent suppression of L1 retrotransposition (Figure 1C and Supplementary Figure S1B and C). Immunoprecipitation experiments demonstrated that HERC5 interacts with ORF1p via the RLD domain (Figure 3A), which is important for reducing the ORF1p level (Figure 2F). Polysome profiling analysis revealed that the L1 RNA enrichment was reduced in heavier polysome fractions, suggesting that HERC5 may decrease L1 translation efficiency (Figure 5E). To our knowledge, this is the first report of translational repression activity of HERC5. Importantly, we did not observe upregulation of global translational-inhibition markers such as phosphorylation of eIF2α and PKR by HERC5 overexpression (data not shown), suggesting that HERC5 acts specifically on L1 translation. Given the enrichment of L1 RNA in the 80S and lighter polysome fractions (Figure 5E), HERC5 may induce ribosome stalling during translation of L1 RNA. Moreover, because HERC5 reduces ORF1p in the RC-L1 RNPs (ORF2p IP fraction) (Figure 5F), we hypothesize that HERC5 binding to L1 RNA also blocks RC-L1 RNP assembly. Further studies will be important to elucidate these mechanisms.

Previous studies have demonstrated that HERC5 inhibits Ebola virus and HIV-1 independently of ISGylation. Although the mechanism by which HERC5 inhibits Ebola virus remains unclear, HERC5 suppresses HIV-1 replication by preventing the nuclear export of unspliced HIV-1 genomic RNA (69). The CRM1/Ran-GTP complex mediates the nuclear export of HIV-1 RNA by interacting with the HIV-1 Rev protein, which binds to the intronic Rev-response element (RRE) within the unspliced HIV-1 RNA. This unspliced HIV-1 RNA export pathway is disrupted by HERC5 in an RLD domain-dependent manner (69, 93, 94). Whether this mechanism affects other unspliced RNAs remains unclear; however, it is less likely to apply to L1 RNA for several reasons. First, HERC5 was initially identified as an L1 inhibitor through our previous proteomics approach, which identified the cytoplasmic L1 RNP complex via ORF1p immunoprecipitation (52). Second, HERC5 is predominantly localized in the cytoplasm, not near the nuclear membrane, where the CRM1/Ran-GTP complex is found (Supplementary Figure S2). Third, we did not observe suppression of proteins translated from unspliced transcripts such as EGFP, luciferase, and monocistronic ORF2, except for ORF1-containing transcripts (Figure 2E, 4B, and Supplementary Figure S4A). Fourth, there is no significant sequence conservation between HIV-1 and L1. Lastly, the identified HERC5 interactors did not include Ran and CRM1 proteins (Supplementary Figure S5 and Table S1). These observations suggest that HERC5 reduces ORF1p expression via a mechanism distinct from the previously reported pathway targeting HIV-1.

Protein expression analyses and *Alu* retrotransposition assays strongly suggested that HERC5 specifically targets the *ORF1* sequence (Figure 2–4 and Supplementary Figure S3). In addition to the L1 RNA-mediated interaction of HERC5 with ORF1p (Figure 3), these results support the notion that HERC5 targets L1 through recognition of the ORF1 RNA sequence. Although HERC5 has been reported to interact with ribosomes and proteins, interactions with specific RNAs have not been described. We considered several possible mechanisms by which HERC5 recognizes the L1 ORF1 RNA. Because HERC5 cannot efficiently inhibit codon-optimized L1 ORFeus retrotransposition, it likely recognizes specific sequences or secondary structures within ORF1 RNA (Figure 3D and E). Additionally, RNA modifications such as m^6^A may serve as recognition markers for HERC5. The number of the canonical m^6^A consensus sequence (5’-RRACH-3’, R = A/G, H = U/A/C) present in the sense strand of *ORF1* varies across elements: 23 motifs in human L1.3, 32 in mouse ORFeus-Mm, and 45 in TG_F_21, but only 16 in human L1 ORFeus. This variation suggests that the inhibitory effect of HERC5 may be reduced in some contexts. Further analysis of target RNAs interacting with HERC5 and motif analysis will be required in the future. Alternatively, HERC5 may be recruited to L1 RNA through interaction with RNA-binding proteins, which were abundantly identified in the proteomics analysis of the HERC5 complex (Supplementary Figure S5 and Table S1). Detailed analyses are needed to determine whether these candidate proteins bridge L1 RNA and HERC5.

In ISGylation, a post-translational modification in which ISG15 is conjugated to substrate proteins (91), not only HERC5 but also other ISGylation components, such as ISG15 and the E2 enzyme UbcH8 (56, 60), are required. Under normal conditions, the expression levels of these proteins are typically low, resulting in limited ISGylation activity. In contrast, ISGylation-independent functions of HERC5 may not require other components and thus can be exerted even in cells without interferon stimulation. In addition to the antiviral effects mentioned, ISGylation-independent roles of hHERC5 and mHERC6 have been implicated in mitochondrial metabolism in non-small cell lung cancer (NSCLC) and in morphogenesis of the seminal vesicle, respectively (95, 96), suggesting that HERC5 and its orthologous proteins are broadly involved in various processes beyond immune responses. Genotype-Tissue Expression (GTEx) data and previous tissue expression analyses show that HERC5 is highly expressed in the testis (97, 53, 98). However, ISG15 expression levels in the testis remain relatively similar to those in other tissues, which suggests that HERC5 also functions independently of ISGylation in the testis (97). Because L1 is highly expressed in germ cells and *de novo* L1 insertions are inherited, multiple pathways suppress L1 in the germline (99–101). The relatively high expression of both L1 and HERC5 in the testis suggests that HERC5 may act as a guardian of the germline genome by restricting *de novo* L1 insertions independently of ISGylation. In addition to previously characterized mechanisms such as RNA degradation (piRNA) and protein degradation (TEX19) (102, 103), a multilayered host defense system, including translational regulation by HERC5, may have evolved in mammals to protect the genome from retrotransposons.

Previous studies proposed a model in which HERC5 interacts with polysomes and mediates ISGylation of nascent peptides during translation (64, 65). We found that the HERC5-polysome interaction and ISGylation functions are independent, suggesting that these activities may have evolved separately. This also supports the idea that polysome interaction and ISGylation were acquired independently in the course of evolution. Among the human small HERC family members examined, only HERC5 suppressed L1 retrotransposition (Figure 6F). Moreover, mHERC6, but not hHERC6, also inhibited L1 retrotransposition, suggesting that mHERC6 acquired the ability after the divergence of mice and humans (Figure 6F). The RLD domains of hHERC5 and mHERC6, especially the first ∼100 amino acids, appear to have undergone positive selection in mammals to gain the ability to interact with polysomes (69, 91, 64, 65). These findings suggest that the small HERC family may have coevolved the capacity to inhibit L1 retrotransposition. Intriguingly, the APOBEC3 gene cluster, comprising ISGs that inhibit both L1 and Alu, is thought to have coevolved with endogenous retroviruses (ERVs) (104, 105). Although evolutionary relationships between viruses and ISGs have been extensively investigated, the coevolutionary dynamics between retrotransposons and ISGs have received much less attention. Future research may further shed light on the evolutionary interplay between these elements.

## Data Availability

Uniprot database (https://www.uniprot.org/help/uniprotkb) was used for protein identification from the mass spectrometry data and the Mascot Server 2.7.0 database (https://www.matrixscience.com) was used as the search engine. The mass spectrometry proteomics data have been deposited to the ProteomeXchange Consortium via the PRIDE partner repository with the dataset identifier PXD068064 and 10.6019/PXD068064 (Token: 4h8ae9WHQpHM). The data supporting the findings of this study are available from the corresponding authors upon reasonable request.

## Supplementary Data statement

Supplementary Data are available at *NAR* Online.

## Acknowledgements

We thank K. H. Burns, H. Siomi, D. Trono, J. L. Goodier, N. Okada, J. V. Moran, A. Roy-Engel, Z. Warkocki, and J. D. Boeke for valuable reagents, all Ishikawa lab members in Kyoto University (especially, K. Onishi and K. Sugino), all RIKEN IMS retrotransposon dynamics team members (H. Suzuki, Y. Fumoto, S. Adachi, K. Yamaguchi, C. Tang, Y. Shinsya, S. Amano, R. Ohara, M. Sato, and E. Kikuchi), S. Azuma and K. Kitao for helpful discussions.

## Author Contributions Statement

K.N., A.L.-F., and T.M. conceived and designed the experiments, analyzed data, and prepared the manuscript. K.N., A.L.-F., and T.M. performed experiments. Y.W. provided technical support, performed mass spectrometry, and contributed to data analysis. M.T. and T.I. provided technical support for ribosome profiling and contributed to experimental design and data analysis. K.N., A.L.-F., F.I., T.I., and T.M. contributed to critical discussions, writing, and editing the manuscript. All authors approved the final version of the manuscript.

