## Supplementary Figures for "The interferon-stimulated gene product HERC5 inhibits human LINE-1 retrotransposition with an ISGylation-independent mechanism"



Table S1 List of interacting proteins shared by HERC5 WT and C994A identified by immunoprecipitation coupled mass spectrometry

Figure S1 HERC5 inhibits L1 retrotransposition independently of ISGylation

Figure S2 HERC5 knockdown increases ORF1p levels; RLD deletion alters HERC5 localization

Figure S3 HERC5 interacts with L1 RNA via the RLD domain

Figure S4 HERC5 targets ORF1p and its downstream ORF2p

Figure S5 Identification of HERC5 WT and C994A common interacting proteins by immunoprecipitation coupled with mass spectrometry

Supplementary Figure S1

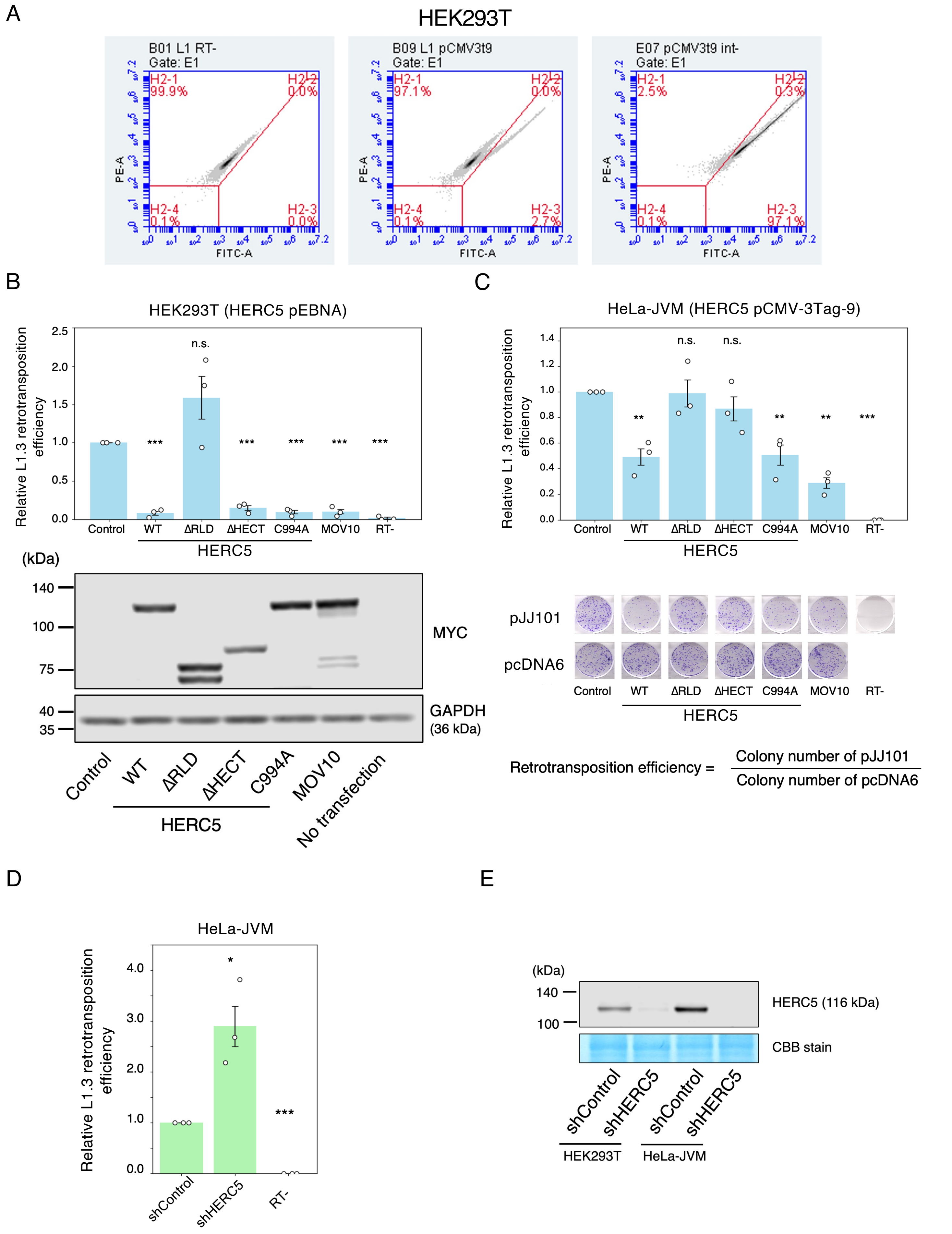

Supplementary Figure S2

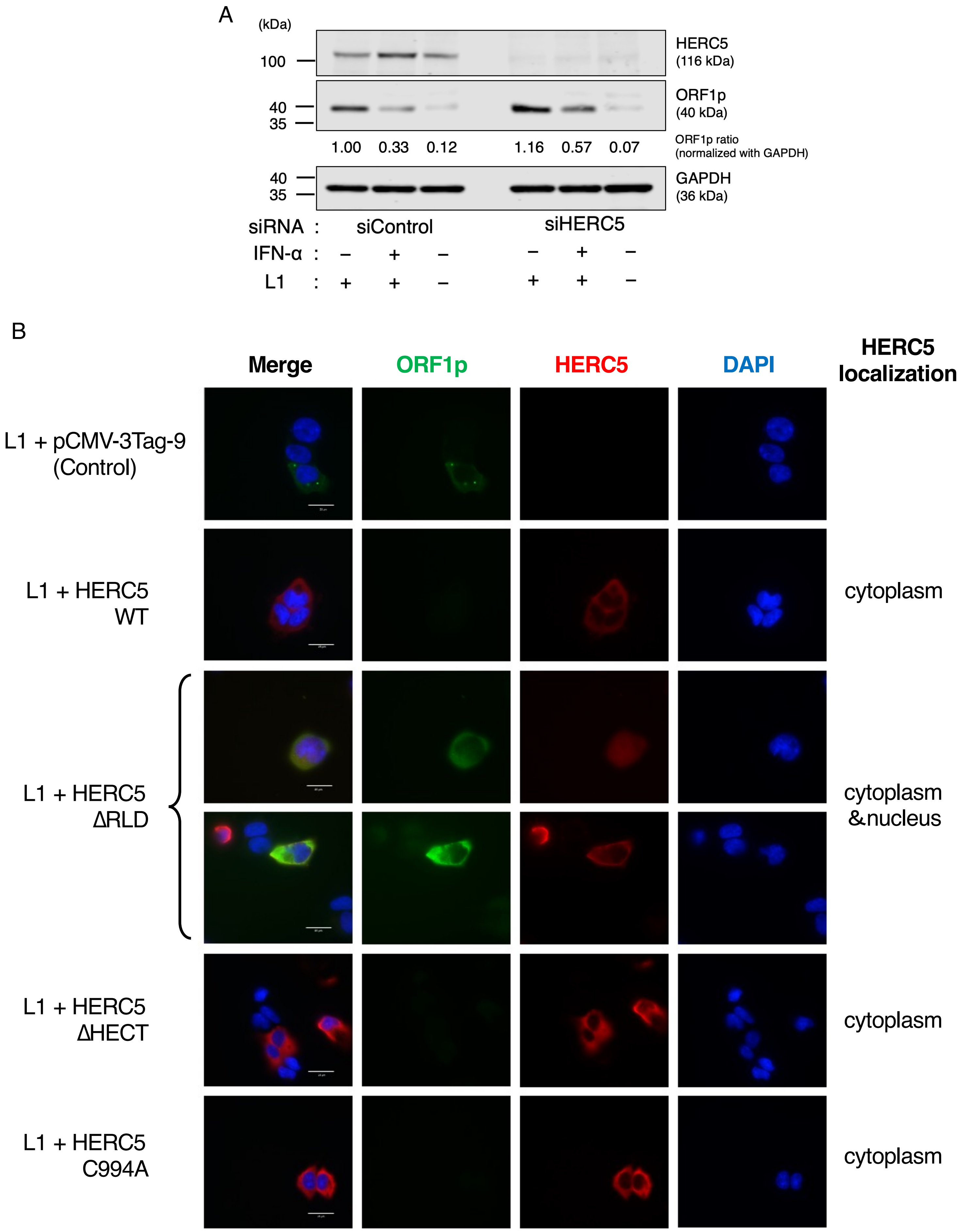

Supplementary Figure S3

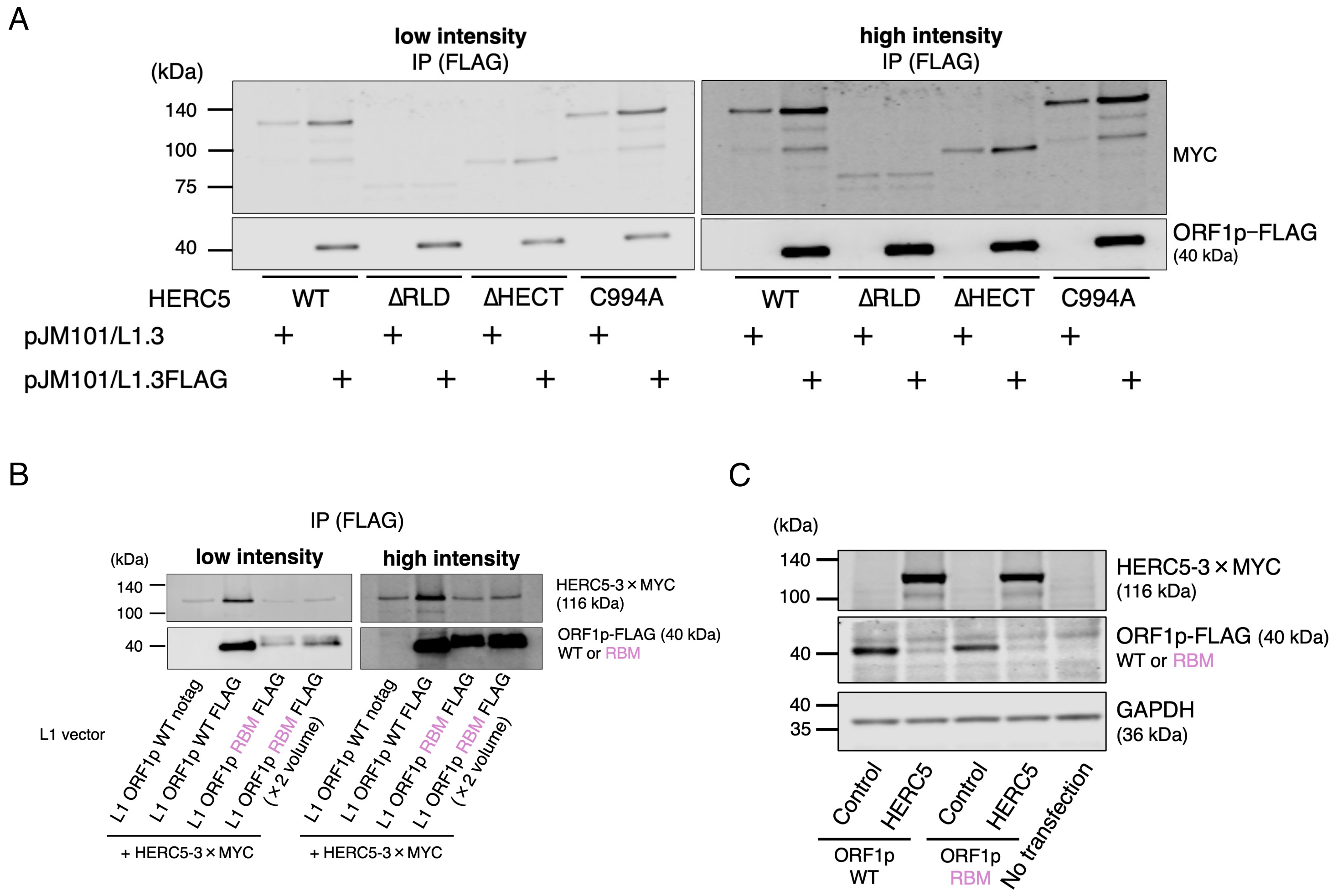

Supplementary Figure S4

A

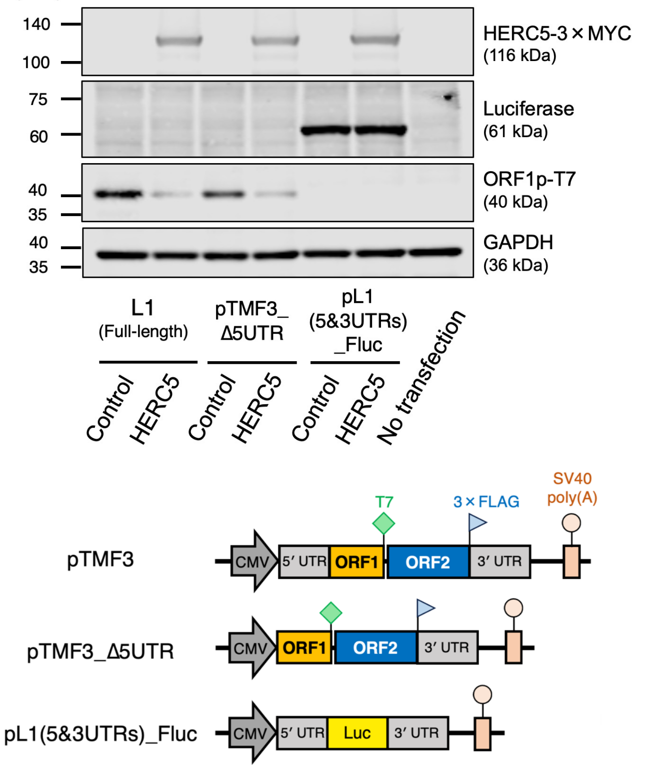


B

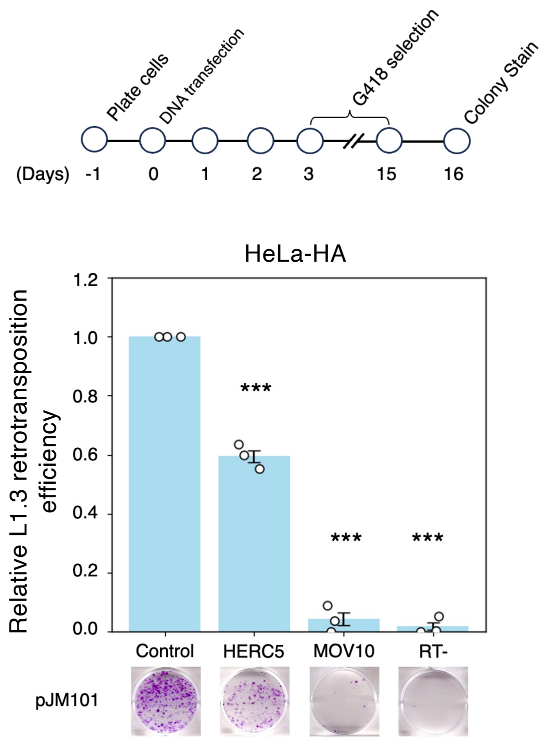

Supplementary Figure S5

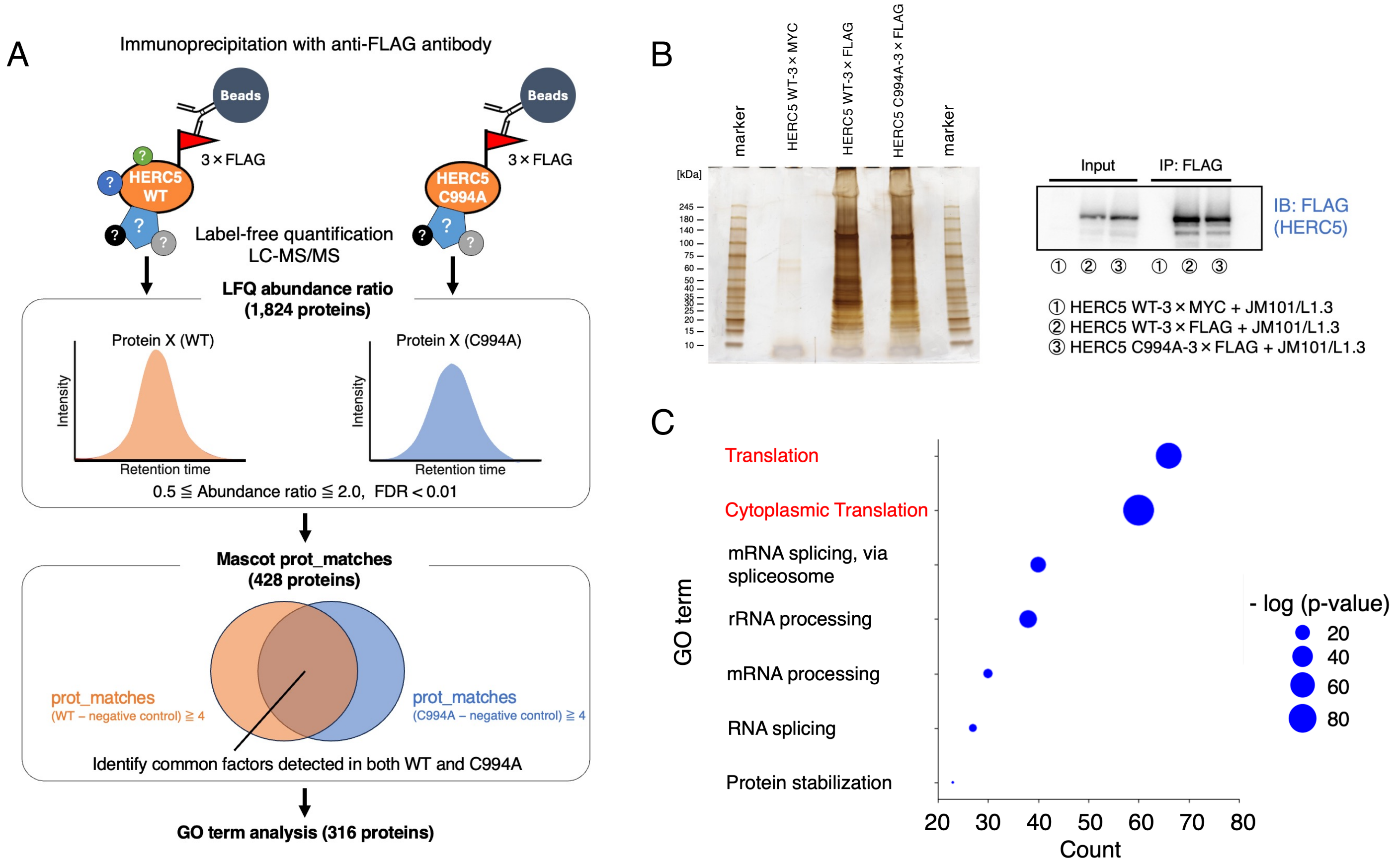

**Supplementary Figure S1. HERC5 inhibits L1 retrotransposition independently of ISGylation**

(**A**) Representative flow cytometry plots of the L1 retrotransposition assay in HEK293T cells using the *mEGFPI* cassette. Left: the threshold line was set based on the scatter plot of cepB-gfp-L1.3RT(-), which served as the negative control. X-axis, fluorescein isothiocyanate-area (FITC-A). Y-axis, phycoerythrin-area (PE-A). Middle: the cells in the H2-3 gate were determined to be EGFP-positive. Right: the percentage of EGFP-positive cells from the intronless EGFP reporter cassette (transfection control). (**B**) L1 retrotransposition assay in HEK293T using the pEBNA vector system. HEK293T cells were co-transfected with an L1-expressing construct (cepB-gfp-L1.3) and either pEBNA (control), HERC5 WT, HERC5 mutants, or MOV10 (positive control). Cells independently co-transfected with cepB-gfp-L1.3RT(-) intronless served as transfection-normalization controls. The transfected cells were selected with blasticidin (10 µg/mL), and the percentage of EGFP-positive cells was determined by flow cytometry. Top: L1 retrotransposition assay with HERC5 overexpression. MOV10 and the RT-deficient L1 (cepB-gfp-L1.3RT[-]) served as controls. X-axis, name of the transfected constructs. Y-axis, relative L1 retrotransposition efficiency compared to the control (pEBNA, set to 1.0). The error bars represent the mean ± the standard error of the mean (SEM) of three independent biological replicates. Each dot represents an independent biological replicate. The *p*-values were calculated using a one-way ANOVA followed by Bonferroni-Holm post-hoc tests; * $p < 0.05$, ** $p < 0.01$, *** $p < 0.001$; n.s.: not significant. Bottom: protein expression of HERC5 WT and mutants expressed from the pEBNA vector system. HERC5 and GAPDH proteins were detected by western blotting using anti-MYC and anti-GAPDH antibodies, respectively. GAPDH served as a loading control. (**C**) L1 retrotransposition assay in HeLa-JVM using the pCMV-3Tag-9 vector system. Top: HeLa-JVM were co-transfected with an L1 expression construct with an *mblastI* retrotransposition indicator cassette (pJJ101) and either HERC5 WT, ∆RLD, ∆HECT, C994A, or MOV10. Cells were selected with blasticidin (10 µg/mL), stained with crystal violet, and the resulting colonies were counted. MOV10 and the RT mutant served as controls. X-axis, name of the transfected constructs. Y-axis, relative L1 retrotransposition efficiency compared to the control (pCMV-3Tag-9, set to 1.0). The error bars and *p*-values were calculated as in (B). Bottom: representative images of stained blasticidin-resistant colonies. The colony numbers of pJJ101 were normalized to those of pcDNA6 to determine retrotransposition efficiency. (**D**) L1 retrotransposition assay with HERC5 knockdown. HeLa-JVM cells were transfected with the WT L1-expressing construct (cepB-gfp-L1.3). The RT-deficient L1 (cepB-gfp-L1.3RT[-]) served as a negative control. After blasticidin selection (10 μg/mL), the percentage of EGFP-positive cells was measured. X-axis, name of the cell line (shControl or shHERC5) or the RT(-) mutant. Y-axis, relative L1 retrotransposition efficiency compared to the control knockdown (set to 1.0). The error bars and *p*-values were calculated as in (B). (**E**) Endogenous HERC5 expression levels in the respective shRNA-knockdown cells. An equal amount of total protein was subjected to western blot analysis. HERC5 was detected using an anti-HERC5 antibody. The membrane was stained with CBB Stain One (Ready To Use) (Nacalai Tesque) and served as a loading control.

**Supplementary Figure S2. HERC5 knockdown increases ORF1p levels; RLD deletion alters HERC5 localization**

(**A**) ORF1p levels with siRNA and IFN-α treatments. HEK293T cells were treated with IFN-α (day −2) and siRNA (day −1), and then transfected with the WT L1-expressing plasmid (pJM101/L1.3). HERC5, ORF1p, and GAPDH were detected using anti-HERC5, anti-ORF1p, and anti-GAPDH antibodies, respectively. GAPDH served as a loading control. ORF1p signal intensities were quantified using Empiria Studio and normalized to GAPDH to obtain the ORF1p/GAPDH ratio. (**B**) Representative immunofluorescence images showing the localization of HERC5 WT, its mutants, and ORF1p. HEK293T cells were co-transfected with pJM101/L1.3FLAG and either the empty vector, HERC5 WT, or the indicated mutants. The localization pattern of HERC5 is noted in the rightmost column. ORF1p was detected in green, HERC5 and its mutants in red and nuclei were counterstained with DAPI (blue). Scale bar, 25 μm.

### Supplementary Figure S3. HERC5 interacts with L1 RNA via the RLD domain

(**A**) Interaction of ORF1p with HERC5 WT or its mutants. HEK293T cells were co-transfected with an L1-expressing vector and a HERC5-expressing vector. ORF1p-FLAG complexes were immunoprecipitated. Left: low-intensity image; right: high-intensity image. HERC5 and its mutants were detected by an anti-MYC antibody, and ORF1p was detected by an anti-FLAG antibody. (**B**) Interaction of ORF1p RBM with HERC5. HEK293T cells were co-transfected with either an L1 WT-expressing vector or an RBM-expressing vector and a HERC5-expressing vector. ORF1p WT and RBM-FLAG complexes were immunoprecipitated. Left: low-intensity image; Right: high-intensity image. ORF1p and HERC5 were detected with anti-FLAG and anti-MYC antibodies, respectively. (**C**) ORF1p WT and RBM levels with HERC5. HERC5, ORF1p, and GAPDH were detected by anti-MYC, anti-FLAG, and anti-GAPDH antibodies, respectively. GAPDH served as a loading control.

**Supplementary Figure S4. HERC5 targets ORF1p and its downstream ORF2p**

(**A**) ORF1p and ORF2p levels with HERC5. Top: HEK293T cells were co-transfected with the modified L1 constructs and a HERC5-expressing vector. The cells were harvested on 3 days post-transfection. HERC5, luciferase, ORF1p, and GAPDH were detected by anti-MYC, anti-luciferase, anti-T7, and anti-GAPDH antibodies, respectively. GAPDH served as a loading control. Bottom: schematic of modified L1 vectors. pTMF3 expresses ORF1p tagged with a T7 gene 10 epitope and ORF2p tagged with a 3×FLAG epitope at their carboxyl termini. pTMF3_Δ5UTR is a derivative of pTMF3 that does not contain the L1 5′ UTR sequence. pL1 (5&3UTRs)_Fluc is a derivative of pTMF3 that contains the firefly luciferase gene in place of the L1.3 coding sequence. (**B**) L1 retrotransposition assay in HeLa-HA. Top: timeline of the assay. HeLa-HA cells were co-transfected with an L1-expressing construct (pJM101) and either pCMV-3Tag-9, HERC5, or MOV10. Cells were selected with G418 (500 μg/mL), stained with crystal violet, and the resulting colonies were counted. The representative images of stained G418-resistant colonies are shown below each condition. The colony numbers of pJM101 were normalized to transfection efficiency to determine retrotransposition efficiency. MOV10 and the RT mutant (pJM105) served as controls. X-axis, name of the transfected constructs. Y-axis, relative L1 retrotransposition efficiency compared to the control (pCMV-3Tag-9, set to 1.0). The error bars represent the mean ± the standard error of the mean (SEM) of three independent biological replicates. Each dot represents an independent biological replicate. The *p*-values were calculated using a one-way ANOVA followed by Bonferroni-Holm post-hoc tests; *** $p < 0.001$.

**Supplementary Figure S5. Identification of HERC5 WT and C994A common interacting proteins by immunoprecipitation coupled with mass spectrometry**

(**A**) Rationale for HERC5 WT and C994A immunoprecipitation-coupled with mass spectrometry. HEK293T cells were co-transfected with an L1-expressing plasmid (pJM101/L1.3) and either HERC5 WT-3×FLAG, HERC5 C994A-3×FLAG, or HERC5 WT-3×MYC (negative control). Cells were harvested on 4 days post-transfection, and HERC5-3×FLAG complexes were purified using an anti-FLAG antibody. Proteins interacting with HERC5 WT and C994A were analyzed by LC–MS/MS, and label-free quantification (LFQ) was performed to determine the relative abundance of proteins. A total of 1,824 proteins were selected based on LFQ abundance ratios and FDR confidence scores, and 428 proteins were selected based on Mascot prot_matches. Gene Ontology (GO) term analysis was performed on the retained 316 proteins common to both filters (abundance ratio and prot_matches criterion) as candidate interactors of HERC5 WT and C994A. (**B**) Silver-stained gel image of proteins co-immunoprecipitated with HERC5. Left: HERC5 WT and C994A complexes purified by immunoprecipitation were separated by SDS-PAGE and visualized by silver staining. The gel was excised into 18 slices according to molecular weight and subjected to mass spectrometry. Right: western blot analysis confirming HERC5 WT and C994A immunoprecipitation. HERC5 was detected using an anti-FLAG antibody. (**C**) Enriched GO terms based on the biological process (BP) among the selected proteins. X-axis, protein count (number of proteins identified by mass spectrometry). Y-axis, name of GO BP terms. Circle size, $-\log_{10}$ (*p*-value). The top enriched GO terms are shown.
